# Clinically relevant humanized mouse models of metastatic prostate cancer to evaluate cancer therapies

**DOI:** 10.1101/2023.10.13.562280

**Authors:** Raymond J. Kostlan, John T. Phoenix, Audris Budreika, Marina G. Ferrari, Neetika Khurana, Jae Eun Cho, Kristin Juckette, Brooke L. McCollum, Russell Moskal, Rahul Mannan, Yuanyuan Qiao, Donald J. Vander Griend, Arul M. Chinnaiyan, Steven Kregel

## Abstract

There is tremendous need for improved prostate cancer (PCa) models. The mouse prostate does not spontaneously form tumors and is anatomically and developmentally different from the human prostate. Engineered mouse models lack the heterogeneity of human cancer and rarely establish metastatic growth. Human xenografts represent an alternative but rely on an immunocompromised host. Accordingly, we generated PCa murine xenograft models with an intact human immune system (huNOG and huNOG-EXL mice) to test whether humanizing tumor-immune interactions would improve modeling of metastatic PCa and the impact of hormonal and immunotherapies. These mice maintain multiple human cell lineages, including functional human T-cells and myeloid cells. In 22Rv1 xenografts, subcutaneous tumor size was not significantly altered across conditions; however, metastasis to secondary sites differed in castrate huNOG vs background-matched immunocompromised mice treated with enzalutamide (enza). VCaP xenograft tumors showed decreases in growth with enza and anti-Programed-Death-1 treatments in huNOG mice, and no effect was seen with treatment in NOG mice. Enza responses in huNOG and NOG mice were distinct and associated with increased T-cells within tumors of enza treated huNOG mice, and increased T-cell activation. In huNOG-EXL mice, which support human myeloid development, there was a strong population of immunosuppressive regulatory T-cells and Myeloid-Derived-Suppressor-Cells (MDSCs), and enza treatment showed no difference in metastasis. Results illustrate, to our knowledge, the first model of human PCa that metastasizes to clinically relevant locations, has an intact human immune system, responds appropriately to standard-of-care hormonal therapies, and can model both an immunosuppressive and checkpoint-inhibition responsive immune microenvironment.

## Introduction

Prostate cancer is one of the leading causes of cancer-related deaths, and treatment options for men with advanced, metastatic disease are limited. Progression to metastatic castration resistant prostate cancer (mCRPC) is often driven by the maintenance of androgen receptor (AR) signaling despite attempted blockades with standard-of-care treatments such as castration and enzalutamide (1,2). Development of novel therapeutics for mCRPC depends on improving the murine models of this disease. The current genetically engineered model (GEM) systems for mCRPC exhibit several shortcomings that hinder their clinical applicability. GEM systems rely on the murine prostate, which differs anatomically and developmentally from the human prostate, and does not form sporadic tumors (3,4). GEMs lack the heterogeneity of human disease and rarely establish metastatic growth, and disease progression in GEM systems tend to be driven in a contrived manner unrelated to human disease or the commonly observed drivers of disease progression (5–7). While human xenografts represent alternative models, they rely on tumor growth in an immunocompromised murine host and are thus unsuitable for investigations into tumor-immune interactions and immunotherapy interventions, a rapidly expanding area of cancer research (5).

Consequently, the development of a human-derived model that can recapitulate the natural history of the disease—from initiation to metastatic spread—and will respond appropriately to the standard of care hormonal therapies is needed to accelerate translational progress in prostate cancer research (5–7). To address this issue, we employed a series of prostate cancer xenograft models in recently developed murine lineages with an intact human immune system (huNOG and huNOG-EXL mice, Taconic Biosciences, Germantown, NY). Male huNOG mice are produced by engrafting juvenile immunocompromised NOG (NOG-NOD/SCID/γnull/c) mice with human CD34+ hematopoietic stem cells (HSC) from human umbilical cord blood. These mice stably develop and maintain multiple human cell lineages, including functional human T-cells, and can be human leukocyte antigen (HLA) matched for a variety of xenograft models (8). Additional mouse models produced with human cytokine [Interleukin-3 (IL-3) and Granulocyte-macrophage colony-stimulating factor (hGM-CSF)] huNOG-EXL, provide more support to human myeloid cells which are often outcompeted by the host in standard huNOG mice (8,9). In the context of enza treatment, we revealed differential levels of metastatic outgrowth in NOG immunocompromised controls compared to newly developed huNOG mice. These results led us to hypothesize that the anti-metastatic responses of AR-targeted therapies are achieved through the human immune system and prompted us to profile the differences in immune microenvironment populations across our models.

## Methods

### Cell lines and culture

R1881 was purchased from Sigma-Aldrich (St. Louis, MO), and enzalutamide (MDV3100), and anti-Programed Death-1 [anti-PD1, Pembrolizumab (Pembro)] were purchased from Selleck Chemicals (Houston, TX), and stored at −20°C in ethanol, −80°C in DMSO, and constituted fresh from lyophilized powder in PBS, respectively. CWR-22Rv1 (22Rv1), and VCaP cell lines were purchased from American Type Culture Collection (Manassas, VA) and were validated and cultured as described (2,10), and transduced with lentivirus for luciferase (luc2) as previously described (2). CWR-R1, VCaP, LAPC4, LNCaP, and enzalutamide-resistant counterparts, in addition to BPH-1, 957E/hTERT, NCI-H660 (H660), PC3, DU145, PNT-2, and RWPE1 cells, were generously provided by Dr. Donald J. Vander Griend at the University of Illinois at Chicago and have been previously characterized and cultured as described (2,11,12). Dr. Peter Nelson at Fred Hutchinson Cancer Center provided LNCaP-sh and LNCaP-APIPC cells (13). All cultures were routinely screened for the absence of mycoplasma contamination using the ATCC Universal Mycoplasma Detection Kit (Manassas, VA).

### Murine prostate tumor xenograft models

NOG control, huNOG and huNOG-EXL humanized mice were obtained from Taconic Biosciences (Germantown, NY). Mice were anesthetized using 2% Isoflurane (inhalation) and either 1×10^6^ VCaP or 5×10^5^ 22Rv1 cells suspended in 100 μl of PBS with 50% Matrigel (BD Biosciences) were implanted subcutaneously into the dorsal flank on both sides of the mice. Once the tumors reached a palpable stage (100 mm^3^), the animals were randomized and treated with enzalutamide or vehicle control [1% Carboxymethylcellulose (Sigma Aldrich), 0.25% TWEEN-80 (Sigma Aldrich), and 98.75% PBS] by oral gavage. Growth in tumor volume was recorded using digital calipers and tumor volumes were estimated using the formula (π/6) (*L* × *W*^2^), where *L* = length of tumor and *W* = width. Loss of body weight during the course of the study was also monitored. At the end of the studies, mice were sacrificed, and tumors were extracted for the downstream analyses. For testosterone implantation, mice were surgically castrated and concurrently implanted with silastic tubing containing 25 mg testosterone (Steraloids Inc, Newport, RI) for sustained release. 22Rv1 cell implantation occurred after 1–1.5 weeks of allowing circulating testosterone levels to equilibrate to approximate human hormone levels (14).

For the VCaP-CRPC experiment, VCaP tumor bearing mice were castrated when the tumors were approximately 200 mm^3^ in size after 14 days. Once the tumor grew back to the pre-castration size, 7 days later, the animals were treated with either vehicle (oral gavage or PBS intraperitoneal injection), enzalutamide (10 mg/kg 5 days a week via oral gavage), pembrolizumab (1 mg/kg 3 times a week, intraperitoneal injection), or both, with respective controls. All procedures involving mice were approved by the University Committee on Use and Care of Animals (UCUCA) at the University of Michigan or Loyola University Chicago and conform to all regulatory standards.

### Ex-vivo imaging

At tumor endpoint, mice were injected with 150 mg/kg body mass D-luciferin (Promega) via intraperitoneal injection, then they were humanely sacrificed, animals necropsied, and tissues were rapidly imaged (within 10 minutes post-sacrifice) with the bioluminescence signal being assessed with the IVIS Spectrum In Vivo Imaging System (PerkinElmer). Tissue tumor burden was calculated based on the total flux (photons per second [p/s]) normalized to area (average radiance), utilizing Living Image software and statistics performed using Graph Pad Prism by utilizing ANOVA with multiple testing corrections across samples and Mann-Whitney and Kolmogorov-Smirnov t-tests between groups.

### Flow Cytometry

#### huNOG experiments

Mononuclear cells were isolated from the subcutaneous tumor and spleen and were stained with fluorescently conjugated antibodies as previously described (15). Quantification of cell number was performed using CountBright Absolute Counting Beads (Thermo Fisher). For cytokine staining, lymphocytes were incubated in culture medium containing PMA (5 ng ml−1), ionomycin (500 ng ml−1), Brefeldin A (1:1,000) and Monensin (1:1,000) at 37 °C for 4 h. Extracellular staining using the antibodies listed below was performed for 20 min, then the cells were washed and resuspended in 1 ml of freshly prepared Fix/Perm solution (BD Biosciences) at 4 °C overnight. After being washed with Perm/Wash buffer (BD Biosciences), the cells were stained with intracellular antibodies listed below. Data collection and analysis was performed on a LSRII equipped with four lasers or a Fortessa equipped with four lasers (BD Bioscience) using BD FACS Diva software. The following human antibodies were used: CD45 (BD Biosciences), CD3 (Thermo Fisher Scientific), CD8 (Biosciences), CD4 (Thermo Fisher Scientific), and IFN-γ (BD Biosciences). All antibodies were used at a 1:100 dilution, as previously described (16).

#### huNOG-EXL experiments

Similar to above, however we utilized the Cytek 5 Laser Aurora ® full spectrum flow cytometer (Cytek Biosciences, Bethesda, MD). Tumor and spleen samples were frozen in liquid nitrogen and later thawed and harvested to acquire mononuclear cells. After harvesting, cells were aliquoted and resuspended in Human TruStainFcX (Clone Information Proprietary, Biolegend) and TruStain FcXPlus (Clone S17011E, Biolegend) according to the provided recommendation (5 ul per 100 ul PBS per 1 million cells). Cells were then washed and stained in L/D Fixable Dead Cell Stain (Thermo Fisher Scientific) for 30 min. Next cells were washed, and extracellular staining was performed using a panel of antibodies listed below for 30 min. Data collection was performed using SpetroFlo 3.0.1. The following antibodies were used: mCD45 (Clone 30-F11, BD Biosciences) hCD45 (Clone HI30, Biolegend), CD3 (Clone UCHT1, Biolgend), CD4 (Clone RPA-T4, Biolegend), CD8a (Clone RPA-T8, Biolegend), CD11b (Clone TCRF44, Biolegend), CD14 (Clone 63D3, Biolegend), CD16 (Clone B13.1, Biolegend), CD19 (Clone HIB19, Biolegend), CD25 (Clone BC96, Biolegend), CD44 (Clone IM7, Biolegend), CD56 (Clone 5.1H11, Biolegend), CD69 (Clone FN50, Biolegend) PD-1 (Clone NA105, Biolegend). Antibodies were titrated for optimal fluorescence per 1 million cells.

### Western blotting

Whole-cell lysates collected from cells seeded at 1×10^6^ cells per well of a 6 well plate (Becton, Dickinson and Company, Franklin Lakes, New Jersey), were lysed in RIPA-PIC buffer [150 mM sodium chloride, 1.0% Igepal CA-630 (Sigma-Aldrich), 0.5% sodium deoxycholate, 0.1% SDS, 50 mM Tris, pH 8.0, 1× protease inhibitor cocktail (Roche Molecular Biochemicals; Penzberg, Germany)], scraped, and sonicated (Fisher Scientific; Hampton, NH; model FB-120 Sonic Dismembrator). Protein was quantified by BCA assay (Thermo-Fisher Scientific), and 50µg of protein were loaded per lane. Antibodies used were: anti-AR (D6F11 XP®, Cell Signaling Technology, Danvers, MA); anti-Beta Actin (AC-15, Sigma-Aldrich); pan-anti-HLA -A-B-C (HLA class I ABC Polyclonal antibody,15240-1-AP, Proteintech group, Rosemont, IL); and anti-PSA (*KLK3*) (D11E1 XP ®, Cell Signaling Technology). Secondary antibodies and Nitrocelluose membranes from Licor (Lincoln, NE) from were used and data captured using a Licor Odyssey M system (Lincoln, NE) as previously described (12).

## Results

### Primary tumor and metastasis assays reveal 22Rv1-engrafted huNOG mice better reflect patient response to castration and enzalutamide treatment than NOG control mice

We first sought to test the growth of the aggressive, bone metastatic CWR-22Rv1 (22Rv1) cell line (2) in huNOG mice compared to NOG [NOG-NOD/SCID/γnull/c (17)] controls with increasing levels of androgen-deprivation via surgical castration and surgical castration with enza treatment (10mg/kg-MDV3100, Selleck Chemical). 22Rv1 are the most aggressive prostate cancer cell line *in vivo* that still maintains AR expression. They express mutant (H875Y), and wild type full length AR and many splice variants including V7, some of which are stably expressed through genetic alterations (18). 22Rv1 respond weakly to both anti-androgens and androgens but are generally considered enzalutamide resistant (19). Isolated from a patient derived xenograft made from a primary tumor from a patient with extensive bone metastases upon disease presentation. In mice, these cells metastasize and colonize clinically relevant sites such as the bone, liver, lungs, and brain (2). We assayed “primary” tumor growth via subcutaneous flank injection (**Figure 1A-B**) and performed metastasis detection and quantification (**Figure 1C-E**, **Figure 2**), with histological validation (representative image in **Figure 1F**), from luciferase-tagged 22Rv1.luc2 [22Rv1 cells transduced with Promega (Madison, WI) luciferase2 ® for bioluminescent imaging] to assay organ-specific metastatic growth (images of the femurs shown in **Figure 1E**, see **Figure 2** for brain, liver and kidneys, **Figure S1** for humerus, **Figure S2** for skull, **Figure S3** for spleen, **Figure S4** for lung, **Figure S5** for heart), of huNOG (two different human male CD34+ HSC donors) and NOG control mice.

**Figure 1:**
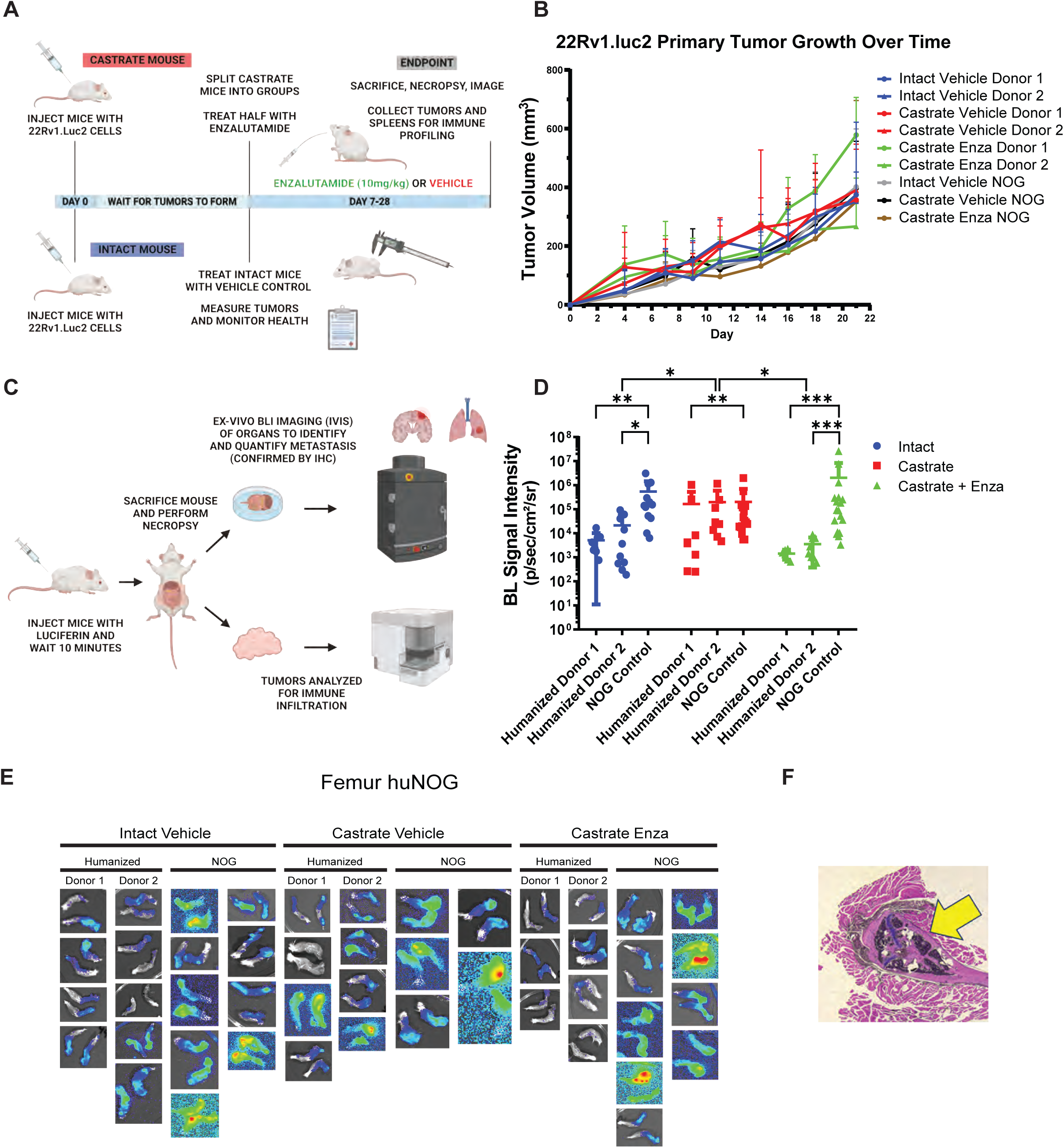
Experimental set-up and 22Rv1 growth in huNOG mice. **A,** Experimental schematic: HuNOG and NOG control mice were surgically castrated. One week following castration, castrated, castrated and enzalutamide treatment and intact control mice were injected subcutaneously with luciferase-transduced 22Rv1 human prostate cancer cells to assay organ-specific metastatic growth. Primary tumors were measured every 2-3 days until endpoint. **B,** Subcutaneous primary flank tumor volume growth measured over time. **C,** Schematic of end-point analysis: Prior to sacrifice mice were injected with luciferin. At sacrifice, organs were ex-vivo analyzed for metastatic growth using the IVIS bioluminescence system PerkinElmer, images taken of signal intensity. **D,** Quantification of average signal intensity per unit area of bioluminescence of mouse femurs. **E,** Representative IVIS images of 22RV1 metastasis to femur. **F,** Histological validation (H&E stain) of the femoral metastases confirmed by a pathologist (Dr. Rahul Manan, MD), with cancer cells seen in both the bone marrow and matrix of the epiphyseal head of a mouse femur (yellow arrow indicates 22Rv1 tumor mass).

**Figure 2:**
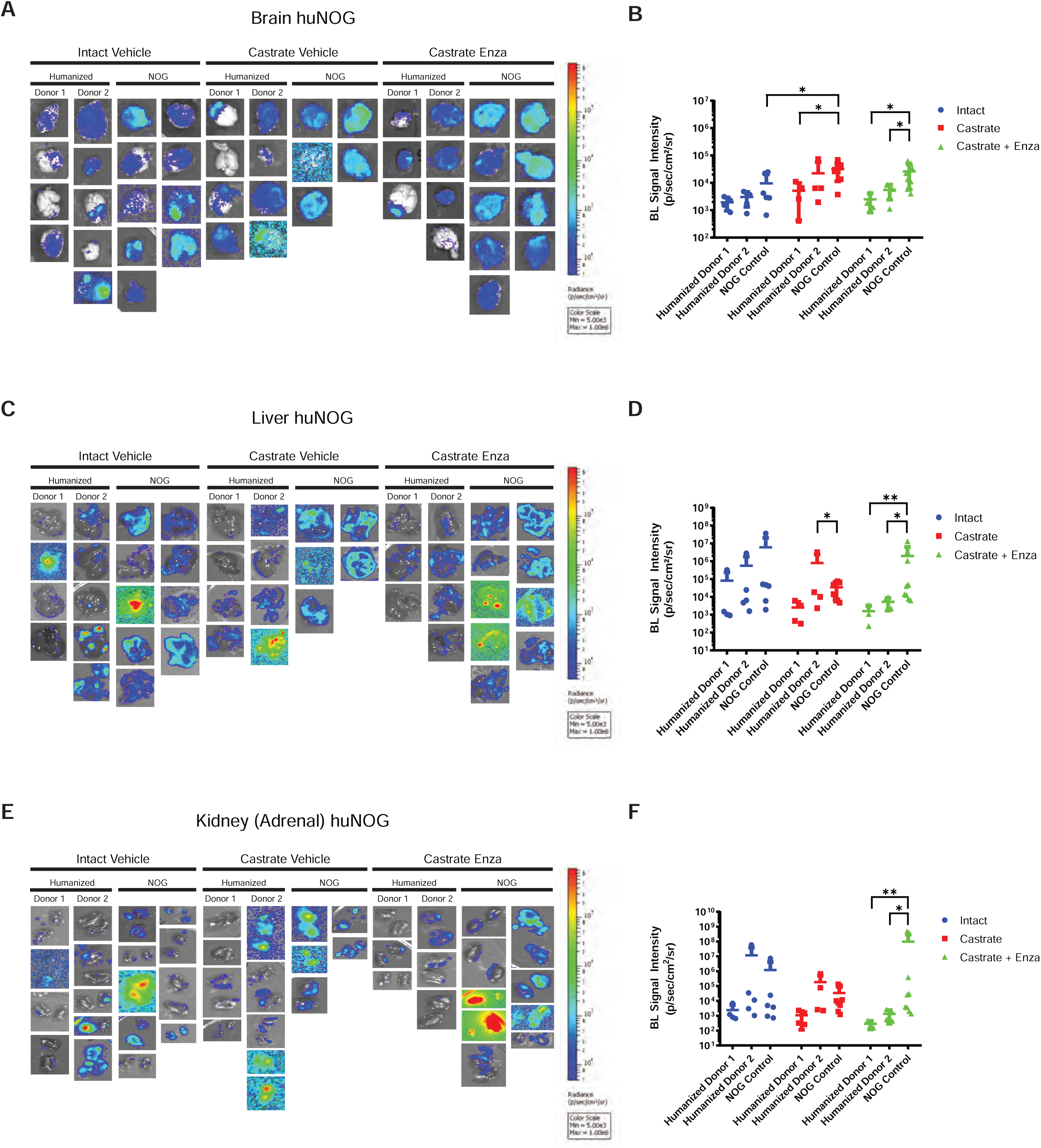
Metastasis by 22RV1 to additional clinically relevant organs in huNOG mice. **A,** Bioluminescent images taken of mouse brain, **B,** Quantified bioluminescence of brain metastasis. **C,** Bioluminescent images taken of mouse liver, **D,** Quantified bioluminescence of liver metastasis. **E,** Kidney (adrenal gland) bioluminescent images, **F,** Quantified bioluminescence of Kidney (tumor mostly in adrenal gland).

The subcutaneous “primary” tumor growth was affected neither by the castration, nor enzalutamide treatment in NOG or huNOG mice at endpoint, but there was variability in presence of an intact immune system in tumor size (**Figure 1B**). At sacrifice, organs were ex-vivo analyzed for metastatic growth using the IVIS bioluminescence system (schematic in **Figure 1C**), and strikingly, metastatic growth was inhibited in huNOG mice under the conditions of castration in combination with enzalutamide treatment, (**Figure 1-2**) mirroring clinical responses which suggest metastatic outgrowth of castration-resistant prostate cancer can be attenuated by enza treatment (20). Surprisingly, AR antagonism with enza in immunocompromised NOG mice, contrary to clinical observations, yet observed in other models (21,22), illustrated increased metastatic colonization and growth (**Figure 1D**). Histology (H&E stain) confirmed femoral metastases (**Figure 1F**), with cancer cells seen in both the bone marrow and matrix of the epiphyseal head of a mouse femur (**Figure 1F**). The effect on growth was also observed for brain (**Figure 2A-B**), liver (**Figure 2C-D**) and kidney/adrenal (**Figure 2E-F**) metastasis in the castration/enzalutamide group for the huNOG mice. These data suggest that 22Rv1 cells have the capacity to metastasize, with or without the presence of an intact human immune system, but the immune system may be activated to prevent outgrowth at secondary sites. These data also suggest that huNOG mice better recapitulate the patient castration/enzalutamide response and warrant further investigation into the immune populations that may be responsible for this phenotype.

### Profiling T-cells in the tumors of huNOG mice reveals an activated immune profile

Since 22Rv1-engrafted huNOG mice exhibited a significant decrease in metastatic colonization compared to control NOG mice (**Figure 1B**), we hypothesized that tumor-infiltrating lymphocytes (TILs) in huNOG mice may be responsible for this suppression, given their anti-tumor effects in patients (23,24). To evaluate TILs in 22Rv1-engrafted huNOG mice, tumors and spleen were collected for immune profiling from treatment groups as described in Figure 1A (either castrate or intact, with the castrate group further divided into enzalutamide treated or untreated). Disassociated tumors from the 22Rv1-engrafted huNOG mice were stained with human α-CD45 (for Leukocytes) and α-CD3+ (T-cells) were co-stained for intracellular IFN-γ (marker of activation), and positively gated for FACS analysis (**Figure 3A**). Analysis of the percentage of CD45+ leukocytes (**Figure 3B**), CD3+ T-Cells (**Figure 3C**), and activated T-Cells (CD3+ IFN-γ+) (**Figure 3D**) showed that there were significant increases in the intra-tumoral CD3+ T-Cells and in their activation state (CD3+ IFN-γ+) in the with enzalutamide treated animals in the castrated group. Overall, the percentage of human CD-45+% immune cells in the tumor was low **(Figure 3B**), and T-cells represented the vast majority of the immune cells in the tumor (**Figure 3C**) in the huNOG model, from this, we focused our analysis solely on the CD3+ population. There were no significant differences in the number splenic of CD4+ helper T-Cells (**Figure 3E**) and CD8+ Cytotoxic T-Cells (**Figure 3F**) for each of the different hormonal and enzalutamide treatment conditions, as a proxy of the whole-body influence of hormones in the mice. However, these data may also be affected by the presence of tumor cells in the spleen (**Figure G**); quantifying splenic metastases, there were variable slight differences across donors and hormone conditions.

**Figure 3:**
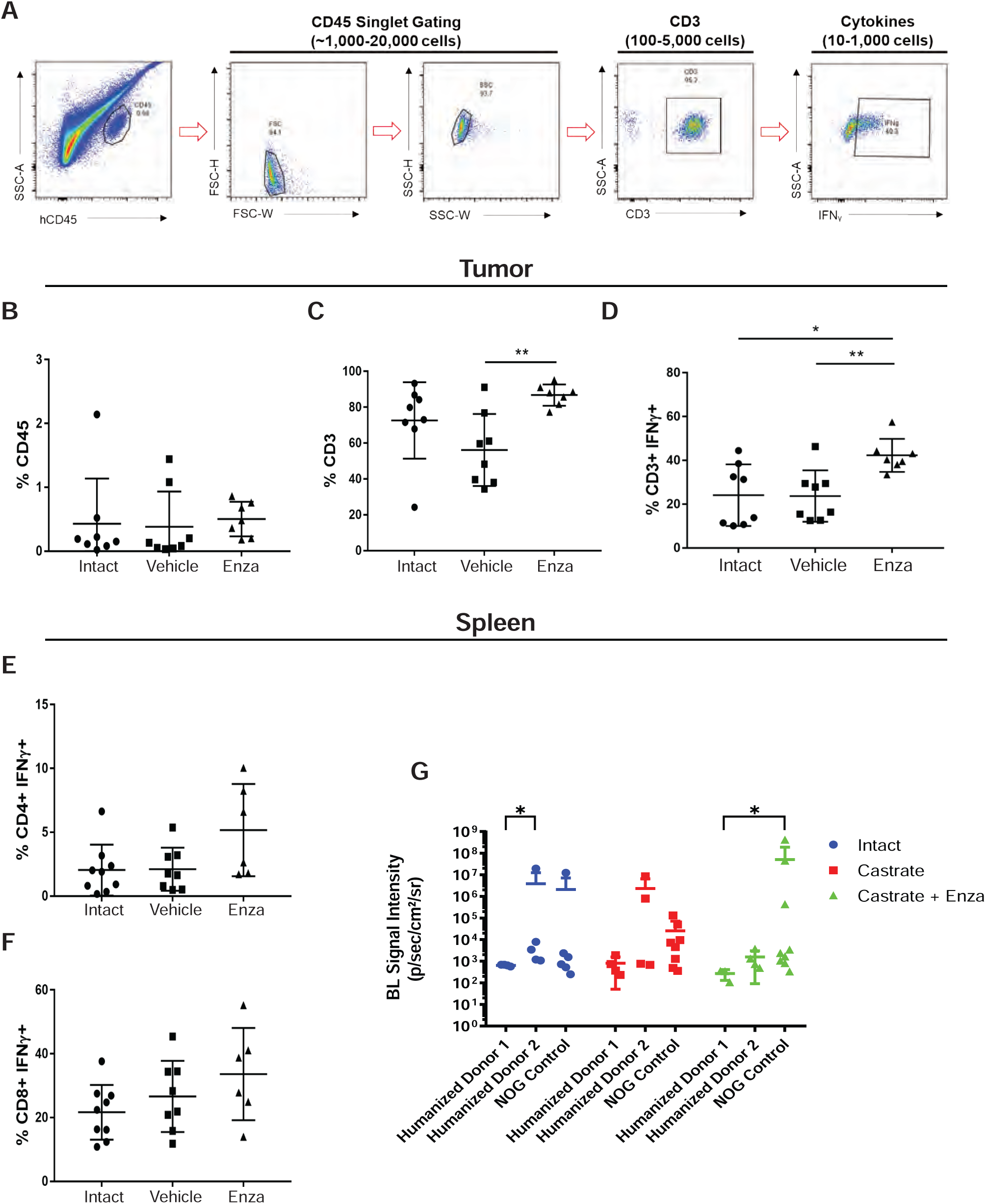
Activated T-cell immune profile in huNOG tumors. **A,** Gating strategy employed to determine T-cell activation status. **B,** Total percentage of CD45+ cells acquired from tumor tissue harvesting split between the intact control, castrated vehicle and castrated with enzalutamide treatment. **C,** Percentage of CD3+ cells observed in the total CD45+ population. **D,** Percentage of CD3+ cells showing IFNg expression via intracellular staining. **E,** Measurement of the percentage of CD4+ cells expressing IFNg harvested from the spleen. **F,** Measurement of CD8+ cells expressing IFNg from the spleen. **G,** Spleen bioluminescent metastasis signal quantification.

### Xenografts in huNOG-EXL mice reveal a suppressed immune profile

Given our results in 22Rv1-engrafted huNOG mice suggested that this mouse model with intact human lymphocytic cells more closely models the patient castration/enzalutamide response than a model using immunocompromised mice; however, immunotherapy responses in prostate cancer patients are low and are often characterized by dense myeloid infiltration (25). We evaluated tumor growth and metastasis in a humanized-NOG mouse system able to maintain not only cells of human lymphocyte lineage, but also human myeloid cells. The huNOG-EXL mice are modified to express human GM-CSF and IL-3 allowing these mice to support human immune cells of myeloid lineage, in addition to lymphocytes. We evaluated subcutaneous tumor growth and measured metastasis in 22Rv1-engrafted huNOG-EXL mice under different levels of androgenic signaling. Mice were divided to four different conditions, listed in descending order of AR-activity outlined in Sedelaar et al. 2013 (26): 1) mice castrated and implanted with testosterone to raise and maintain testosterone levels (530 ± 50 ng/dL) that are physiologically relevant in normal humans (26); 2) mice with intact gonads, thus mimicking hypogonadal or castrate human levels; 3) mice castrated, thus mimicking patients treated with abiraterone (the CYP17A inhibitor (27)); 4) mice castrated and dosed with enzalutamide, thus mimicking androgens depleted and AR-antagonized conditions. Similar to the result in 22Rv1-engrafted huNOG mice with two different HSC donors, primary tumor growth was not affected by either castration or enzalutamide treatment (**Figure S6**). But unlike 22Rv1-engrafted huNOG, no difference in metastasis was observed in the huNOG-EXL mice treated with enzalutamide (**Figure 4A-H**) suggesting a possible role of mature myeloid cells in preventing the enzalutamide effect on metastatic growth seen in huNOG mice. This suggests the possibility of immunosuppressive myeloid cells in tumors of huNOG-EXL mice.

**Figure 4:**
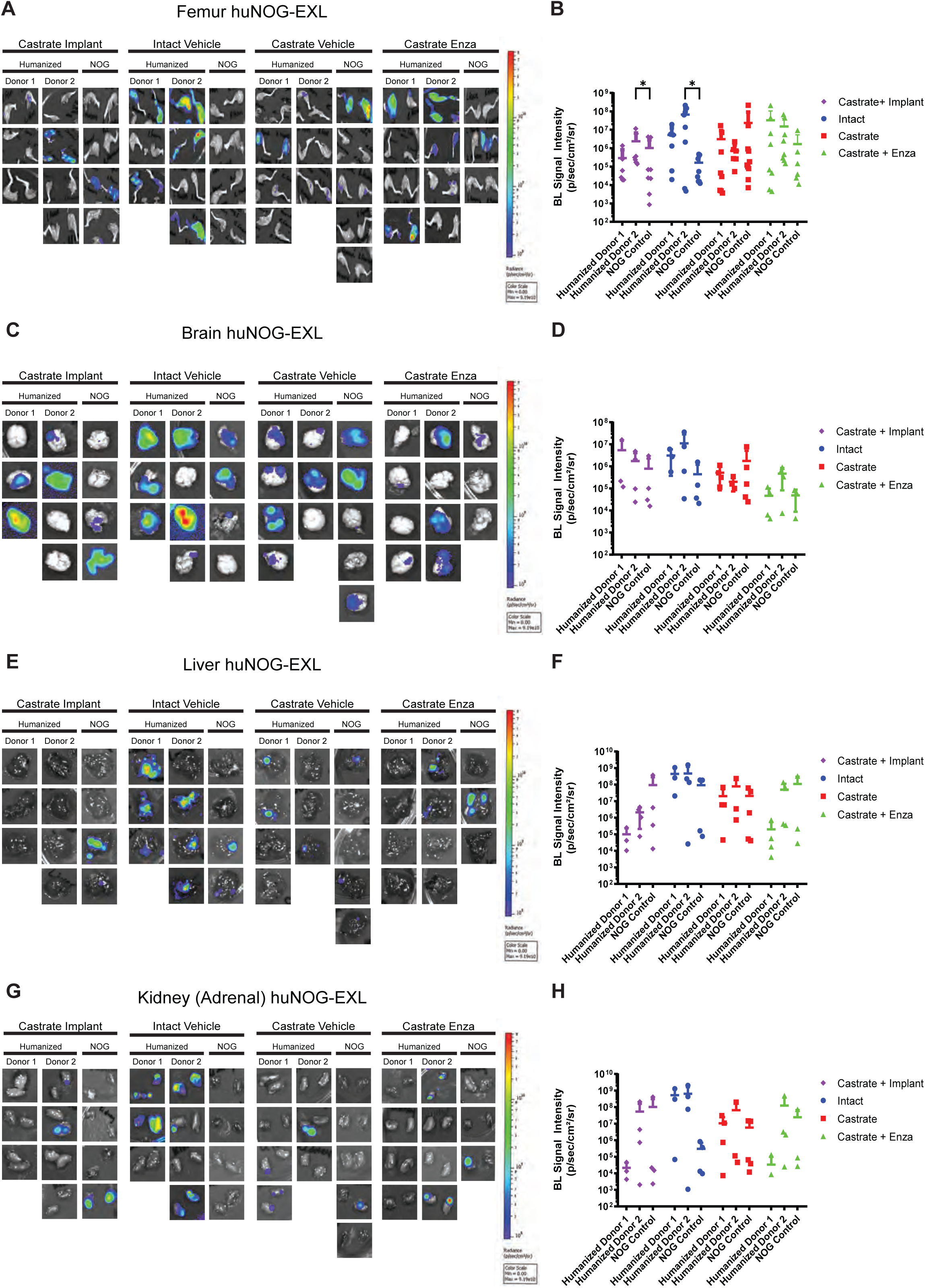
Metastasis by 22RV1 to additional clinically relevant organs in huNOG-EXL mice. **A,** Bioluminescent images taken of femurs, **B,** Quantified bioluminescence of femur metastasis. **C,** Bioluminescent images taken of mouse brain, **D,** Quantified bioluminescence of brain metastasis. **E,** Bioluminescent images taken of mouse liver, **F,** Quantified bioluminescence of liver metastasis. **G,** Kidney (adrenal gland) bioluminescent images, **H,** Quantified bioluminescence of Kidney (tumor mostly in adrenal gland).

Our data from TILs (**Figure 3**) showed that there are activated T-cells in the tumors of 22Rv1-engrafted huNOG mice. We immunologically profiled subcutaneous tumors from 22Rv1-engrafted huNOG-EXL mice using the Cytek Aurora ® system to understand the difference in enzalutamide effect in huNOG compared to huNOG-EXL mice (See details of antibody panel in methods). TILs from 22Rv1-engrafted huNOG-EXL mice tumors were collected for immune profiling from treatment groups with descending levels of androgenic signaling as described for **Figure 4** [either castrated/testosterone implant (Test), intact/vehicle (Int), castrated/vehicle (Cast) or castrated/enzalutamide (Enza)]. Analysis of the population of leukocytes (CD45+), T-Cells (CD3+), helper T-Cells (CD4+) and myeloid cells (CD3-CD19-CD11b+) (**Figure 5A**) showed no differences between the treatments in the intra-tumoral populations of these immune cells (**Figure 5B-C**) nor any major differences within the spleens (**Figures S7-S9**) in the huNOG-EXL model, with similar levels of human leukocyte percentages in the overall tumor. The populations, and activation levels determined by CD25 [the IL-2 receptor, induced upon initial T-cell activation, constitutively expressed in T-regulatory cells (Tregs) (28)], CD44 [upregulated upon activation in effector and memory T-cells (29)], CD69 [a broadly expressed early activation marker in leukocytes implicating in retaining cells in peripheral tissues (30)] and PD1 [programmed cell death-1, a marker of T-cell exhaustion and Treg differentiation (31)] abundance (**Figure 5D**), of CD14 negative myeloid cells also showed no differences between the treatments in this model (**Figure 5E-F**). The activation level was low, suggesting the huNOG-EXL represent an immune “cold” model similar to what is seen in the majority of prostate cancer patients (32,33). The population of CD14+ myeloid cells observed was too small to determine their level of activation (**Figure 5**). In fact, it is interesting to note that for all treatments, the majority of myeloid cells were CD14-negative, which is indicative of immature myeloid derived suppressor cells (MDSCs) (34). We identified very few tumor-infiltrating B-cells (CD19 positive) and Natural Killer (NK) cells (CD56+, CD16+) across hormone conditions, despite high levels in the spleens of these mice (**Figure S9 E-G**).

**Figure 5:**
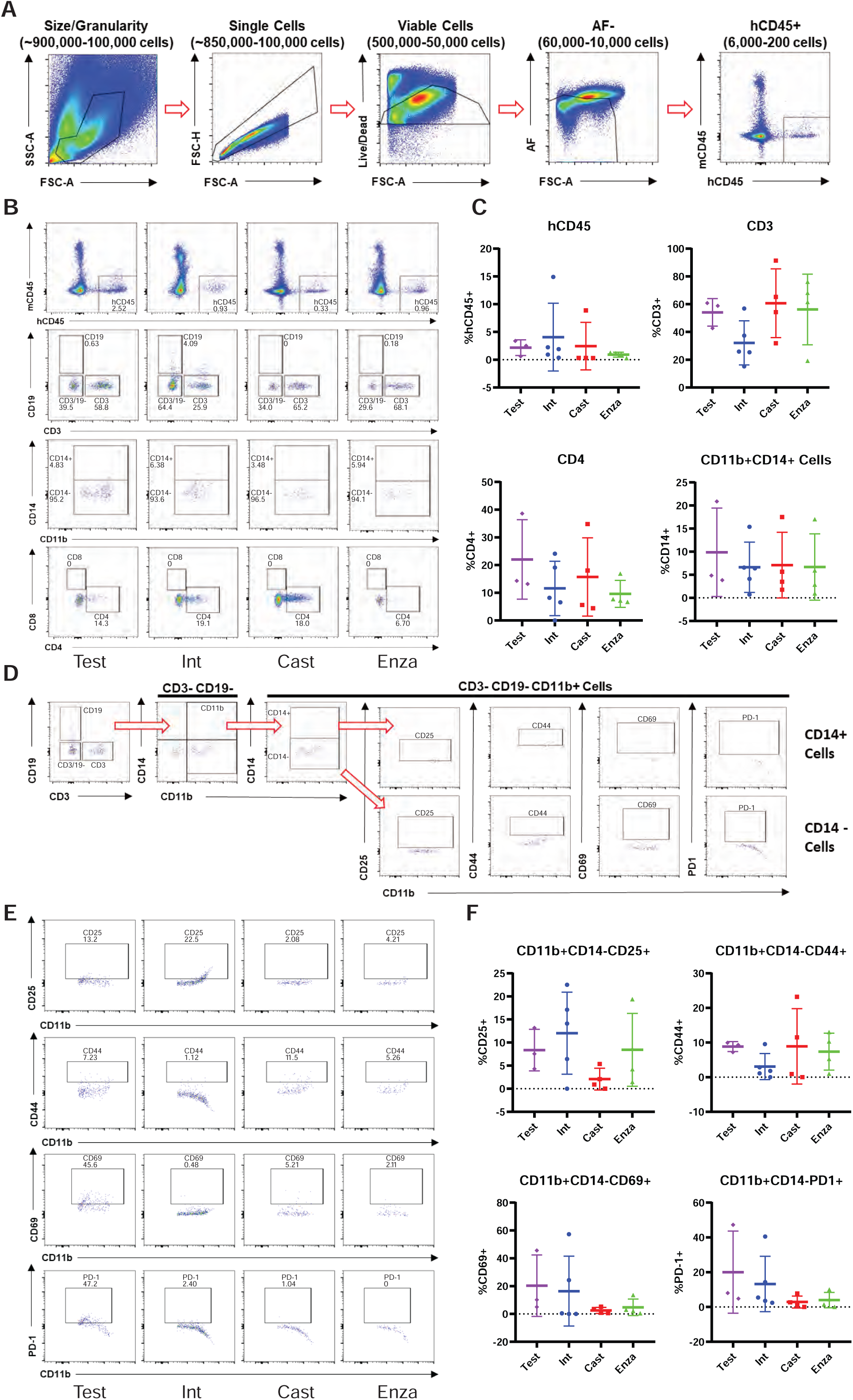
Myeloid-support exhibits immuno-dampened profile in huNOG-EXL 22Rv1 xenograft tumors. **A,** Gating strategy used to determine presence of human CD45+ cells in the NOG-EXL model. **B,** Representitave data showing the abundance of various immune cell populations; human leukocytes, CD19+ cells, CD3+ cells and double negative cells, MDSC and activated myeloid cells and helper t-cells (CD4+) and cytotoxic t-cells (CD8+) (top to bottom). **C,** Quantitated data comparing human CD45, CD3, CD4, and CD11b populations in tumors isolated from testosterone implanted vehicle, intact vehicle, castrated vehicle, and castrated enzalutamide treated mice. **D,** Gating strategy for determining the activation state of MDSCs (CD3-CD19-CD11b+ CD14-). **E,** Representative data showing the activation state of the MDSC cells harvested from tumors under different treatment categories through the presence of the surface markers: CD25, CD44, CD69 and PD-1. **F,** Quantitated data showing the activation state of the MDSCs throughout the different treatments.

We also profiled T-cells from 22Rv1-engrafted huNOG-EXL mice tumors for activation markers and to determine the population of Tregs (**Figure 6A**). Although no differences were observed between treatment groups, of the CD3+ CD25+ cells, the majority were PD1+ which is indicative of the regulatory T-cell phenotype, and found in the bone-microenvironment of prostate cancer patients (16). This result, together with the data on MDSCs (**Figure 5**) suggests a suppressed immune profile (“cold”) in the huNOG-EXL system, in contrast to the activated immune profile (“hot”) seen in the huNOG model (**Figure 3**).

**Figure 6:**
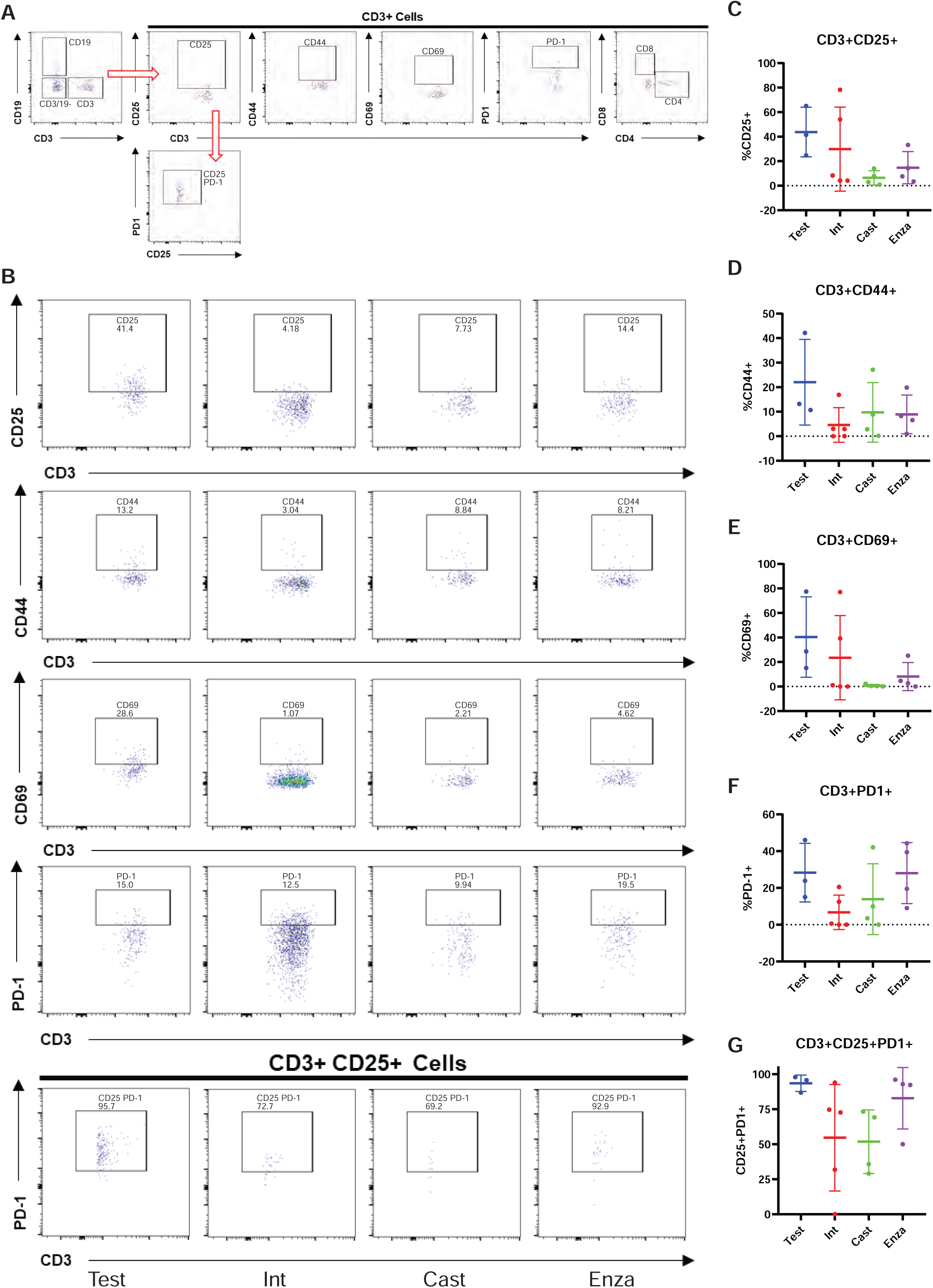
Immune-dampened profile of t-cells in huNOG-EXL tumors. **A,** Gating strategy for determining the activation state of t-cells and regulatory-like t-cells. **B,** Representative data showing the activation state of the CD3+ cells harvested from tumors under different treatment categories through the presence of the surface markers: CD25, CD44, CD69 and PD-1 and regulatory-like cells (CD3+CD25+PD-1+). **C,** Quantitation showing the expression of CD25, **D,** CD44, **E,** CD69, **F,** PD-1, **G,** regulatory-like t-cells.

### VCaP Xenograft Tumors respond to Immunotherapy and Enzalutamide in huNOG Model

To test another xenograft model in humanized mice, we relied on the VCaP castration-resistant-xenograft model, which mimics the natural history of prostate cancer, with short-term responses to both castration and AR-antagonism (19). We assayed primary tumor growth using a VCaP castration resistant model in huNOG mice to evaluate the model in a more enzalutamide sensitive setting, and given our results with 22Rv1, determine if we could enhance the effects of immune activation by treatment of a checkpoint inhibitor in the form of the anti-PD1 pembrolizumab (pembro). In this model VCaP tumor–bearing mice were castrated when the tumors were approximately 200 mm^3^ in size, and once the tumor grew back, animals were randomized and treated with enzalutamide and/or the anti-PD-1 antibody pembrolizumab (pembro), or with vehicle controls. VCaP xenograft tumors treated with enza and pembro work well as monotherapies, and slightly better in combination, completely eliminating tumors. (**Figure 7A**). NOG mice were not responsive to either of these treatments (**Figure 7B**). This result demonstrates that this model is responsive to immune checkpoint inhibition and works synergistically to AR targeted therapy.

**Figure 7:**
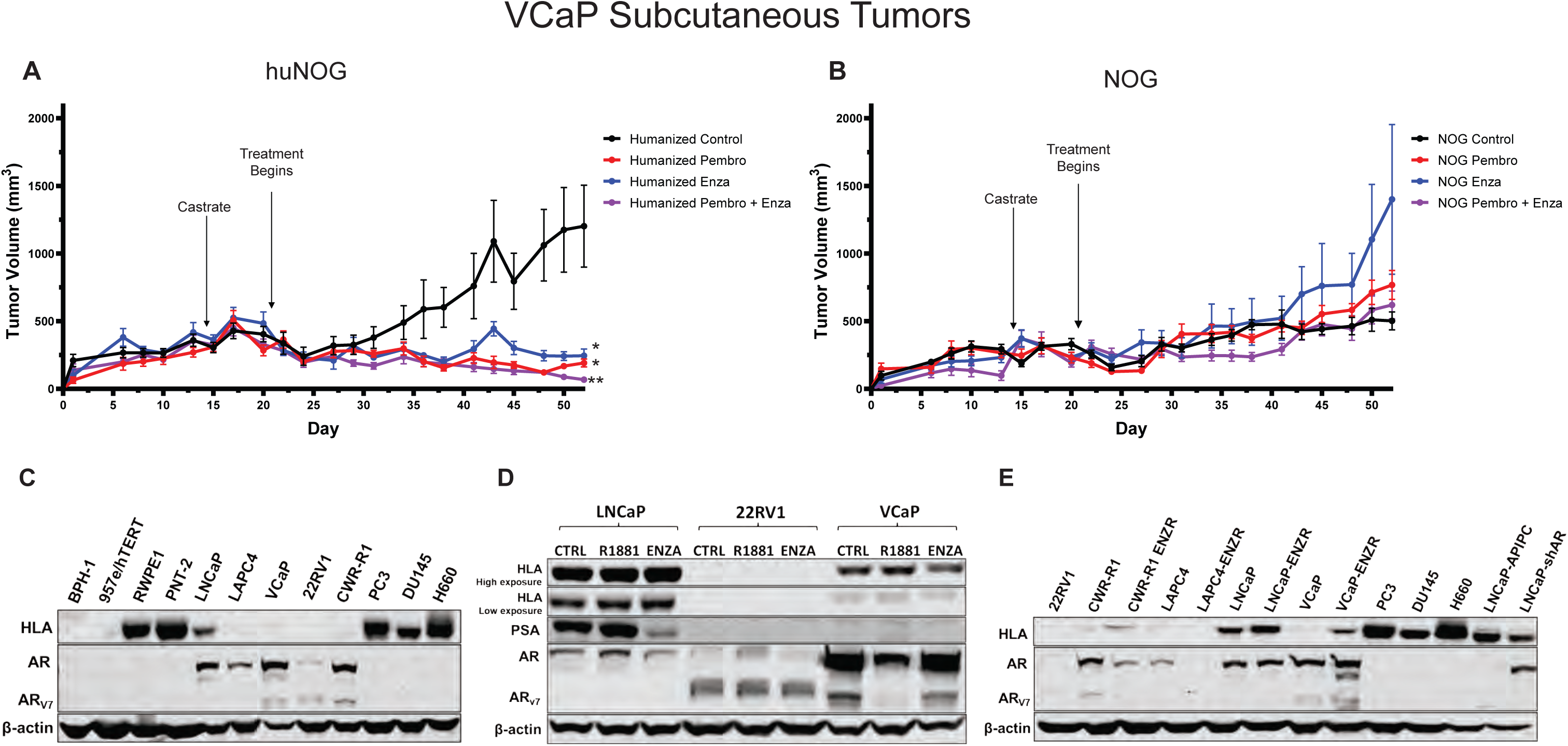
Response of VCaP subcutaneous tumors to anti-PD1 immunotherapy and enzalutamide. **A,** huNOG primary tumor growth response and **B,** NOG primary growth response to pembrolizumab, enzalutamide and combination therapy. * Represents p>0.05 when compared to vehicle control, ** represents p>0.05 when compared to all other conditions. **C,** HLA (HLA – A, -B, -C) expression of a panel of benign prostate (BPH-1, 957e/hTERT, RWPE1, PNT-2), AR-positive prostate cancer (LNCaP, LAPC-4, VCaP, 22Rv1, CWR-R1) and AR-negative (PC3, DU145 and NCI-H660) cancer cell lines determined by western blotting (representative blots show, experiment replicated three times). Corresponding AR blot and AR variant (AR-V7) detection. **D,** LNCaP, 22RV1 and VCaP HLA response the AR agonist, R1881 (1 nM), AR-antagonist enzalutamide (ENZA - 10 μM) and vehicle control (CTRL) treatment for 24 hours. PSA is a canonical AR-target gene as a readout of AR modulation. **E,** Comparison of HLA expression between enzalutamide naïve and resistant cell lines (ENZR).

Finally, to provide some insight of the mechanism of these effects seen in the huNOG mice, we assayed prostate cancer cells for HLA expression (**Figure 7C**) and to identify Androgen-control of HLA (-A, -B, -C) expression. 22Rv1 cells display no expression of HLA molecules, regardless of AR modulation (**Figure 7C, D, E)**, however, VCaP cells illustrated increased HLA expression in the enzalutamide-resistance state (**Figure 7D**), and slight decreases with short-term treatment (**Figure E**). Other AR-positive prostate cancer cell lines generally display decreases in HLA expression when compared to immortalized benign controls (**Figure 7C)**, and cells that maintain AR-post enza-resistance show increased HLA expression, as evidenced by previously reports (2,33,35,36). Taken together, these data suggest that much of this response is likely governed by the T-cell activation seen with enza-treatment (37), as well as some potential cancer cell autonomous effects, all of which are topics of future investigation.

## Discussion

These results illustrate, to the best of our knowledge, the first model of human PCa that metastasizes to clinically relevant locations, has intact human immune system, and responds appropriately to standard-of-care hormonal therapies. We found that humanizing tumor-immune interactions improved modeling of metastatic PCa and provides a model more suitable to evaluate hormonal and immunotherapies. Use of huNOG mice provided a model that not only has an intact human immune system, but also shows metastases to relevant secondary sites and models the effect of anti-androgen treatment on metastasis.

The 22Rv1-engrafted huNOG mouse models human PCa in that it metastasizes to clinically relevant locations. Metastasis was observed in bone (femur and humerus), lymph nodes, liver, brain, spleen, lungs, and kidneys (adrenal) in 22Rv1-engrafted huNOG as well as in the immunocompromised NOG mice. However, reduction of metastasis by enzalutamide was only observed in the huNOG mice. In the immunocompromised NOG mice enzalutamide treatment did not decrease metastasis, but instead showed a paradoxical increase in metastases. The decreased metastasis with enzalutamide treatment in the huNOG is consistent with the clinical situation where Hussain *et al*. found that enzalutamide treatment of patients with non-metastatic CRPC had a 71% lower risk of metastasis (20). In this way, the xenograft model in huNOG mice has the advantage that it responds appropriately to standard-of-care hormonal therapies.

An important feature of this xenograft model in huNOG mice is the presence of tumor-immune interactions. The huNOG mice maintain cells of the human immune system, including functioning human T-cells. Taken together, the results suggest that the effect of enzalutamide to decrease metastasis involves an interaction with the immune system. This hypothesis is supported by the presence of increased infiltration of activated (INFγ+) T-cells into the tumors from the huNOG mice treated with enzalutamide. As androgen signaling has been shown to be immunosuppressive (37,38), AR-antagonism might relieve an inhibitory signal and activate immune surveillance, as well as promote T-cell tumor infiltration. These results suggest that a huNOG xenograft model exhibits an activated “hot” immune profile from an immune perspective and this aspect may be important in the ability of the model to replicate the ability of enzalutamide to decrease metastasis as seen in the clinical situation.

Interestingly, the enzalutamide effect to decrease metastasis was not observed in the huNOG-EXL model. The huNOG-EXL mice maintain human immune cells of both a myeloid and lymphoid lineage. The TILs from tumors from enzalutamide treated huNOG-EXL mice did not show the increased infiltration of activated T-cells that was observed with the huNOG mice, but for all treatments in the huNOG-EXL, the presence of MDSCs and Tregs suggests a suppressed “cold” immune profile in the huNOG-EXL system. This immunosuppressed profile may account for the lack of effect of enzalutamide on metastasis in this system. Enzalutamide itself has been shown to promote immunosuppressive effects in myeloid populations through a non-AR dependent manner (39).

The huNOG mouse xenograft model can be used for other PCa tumor cell lines, besides 22Rv1. Using a VCaP castration resistant model in huNOG mice, we demonstrated the usefulness of a huNOG mouse xenograft to evaluate immunotherapies, either as monotherapies or in combination with a hormone treatment such as enzalutamide. VCaP xenograft tumors were responsive to the anti-PD1 pembrolizumab as a monotherapy, and in combination with enzalutamide worked synergically (**Figure 7A**). Tumors in immunocompromised NOG mice were not responsive to either of these treatments (**Figure 7B**).

Future work will be focused dissecting the mechanism of how AR and its signaling exhibit the cell-specific effects within prostate tumors, and how this affects interactions between cancer cells and immune cells in the tumor microenvironment to produce differential responses to AR targeted therapies. We observed that enzalutamide resistant cells upregulate HLA, here (**Figure 7E**) and as previously reported (2), but others have also illustrated a connection between chronic enzalutamide treatment and upregulation of immune-regulatory proteins that produce an immunosuppressed microenvironment (40). Our models presented here have the potential to aid further work which is necessary to tease apart the timing and phasing of AR inhibition to produce an optimal immune response. Responses excitingly illustrated for a subset of patients with the use of as biphasic androgen therapy has produced responses to AR-checkpoint inhibition (41). Finally, we illustrate the use of this model across a few cancer cell line derived xenografts; however, we hope to utilize mice with humanized immune systems to model the diverse clinical responses seen to AR-and immune-checkpoint inhibition in other tumor models. Particularly those from patient derived xenografts and organoids, such as CDK12 mutated tumors – some of which are responsive to checkpoint inhibition (42) – and to model clinical resistance to different therapies and disease states that otherwise has been difficult to do with standard xenografts or GEM models.

## Acknowledgements

We would like to thank the Prostate Cancer Foundation for their generous support and guidance for this project, particularly Drs. Howard Soule, PhD and Andrea K. Miyahira, PhD. We would like to thank Taconic Biosciences, particularly Dr. Paul Volden, PhD, as well as Drs. Philip Dube, PhD and Michael Seiler, PhD for their support and expertise. We would also like to acknowledge Stephanie Simko, Parth Desai, Xuhong Cao, Jean Tien, PhD, and other members of the Vander Griend and Chinnaiyan labs for their technical support. We would like to thank Dr. Peter Nelson for the LNCaP-sh and APIPC cell lines. We would also really like to thank the Department of Cancer Biology at Loyola University Chicago and Department of Pathology at University of Illinois at Chicago, and the support of their respective chairs, Dr. Nancy Zeleznik-Le, PhD, and Dr. Alan Diamond, PhD. Finally, we would like to thank the Cardinal Bernardin Cancer Center FACs Core Facility and the Cellular Therapy Center Core Facility, director Dr. Phong Le, PhD, and the help received to us by Pat Simms, Bert Ladd, and Corbin Pomykata. S.K. was supported by a Cancer Biology Training Grant (T32-CA09676), the Department of Defense (DoD) Prostate Cancer Research Program (PCRP) Early Investigator Research Award (W81XWH-17-1-0155), and the 2019 Robert Citrone, George Walker, & Mark Weinberger – Prostate Cancer Foundation VAlor Young Investigator Award. S.W. is supported by NCI R01 (R01CA215758). A.M.C. is an NCI Outstanding Investigator, United States (R35CA231996), Howard Hughes Medical Institute Investigator, United States, A. Alfred Taubman Scholar, and American Cancer Society Professor, and is supported by NCI R01 (R01CA200660), United States.

---

**Figure S1:**
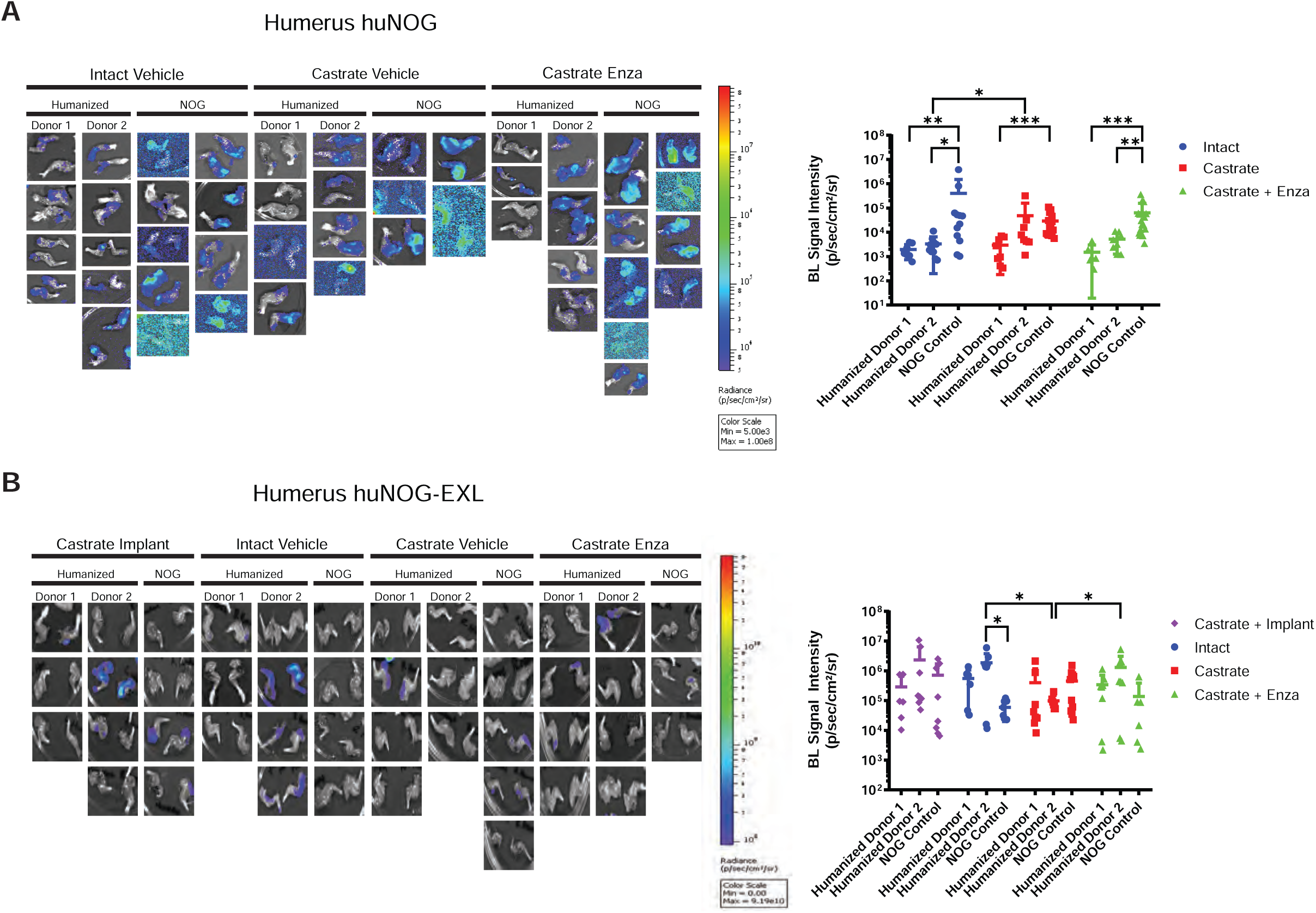
Metastasis by 22RV1 to the humerus in huNOG and huNOG-EXL mice. **A,** Bioluminescent images taken of huNOG humeri, **B,** Quantified bioluminescence of huNOG humerus metastasis. **C,** Bioluminescent images taken of huNOG-EXL humeri, **D,** Quantified bioluminescence of huNOG humerus metastasis.

**Figure S2:**
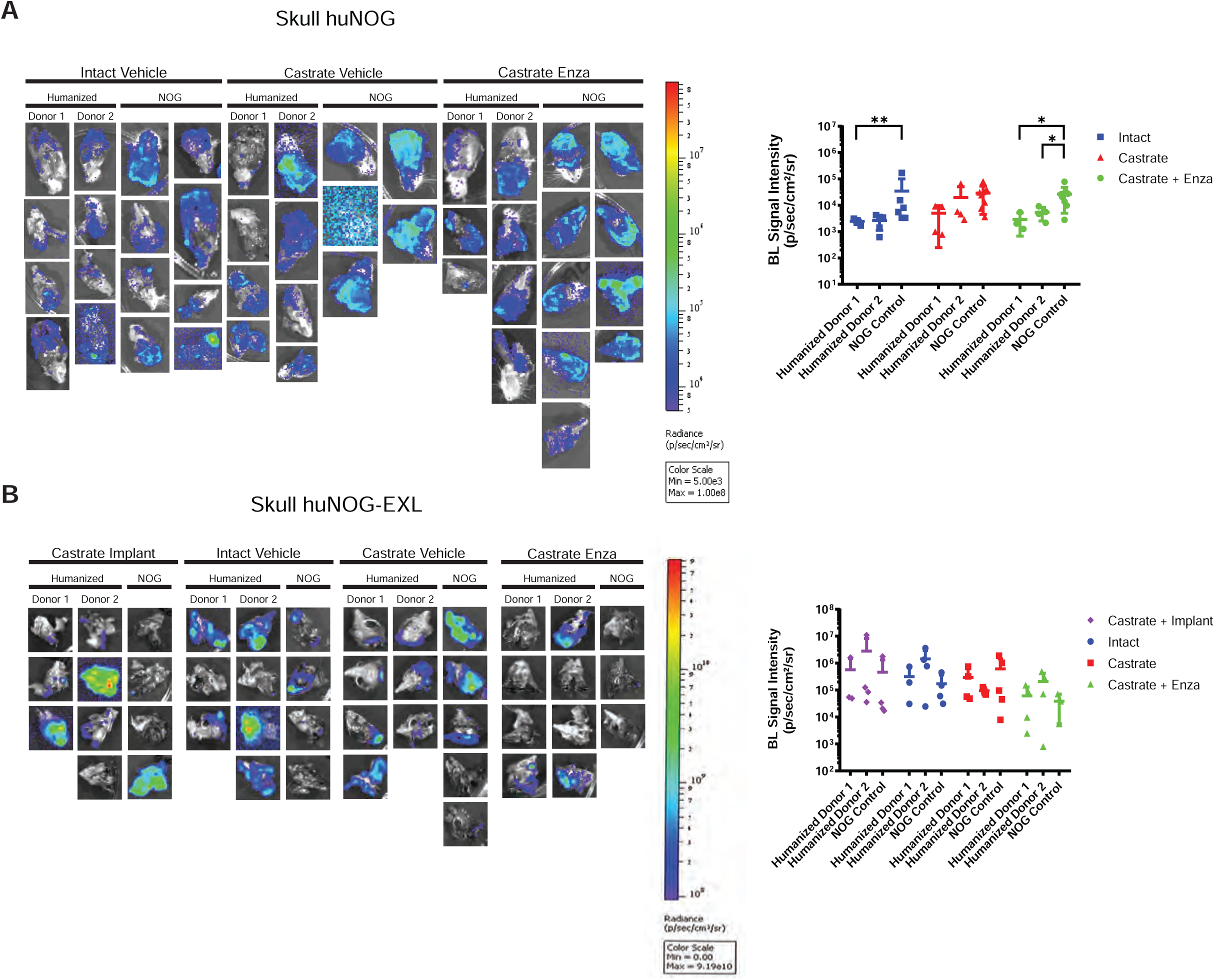
Metastasis by 22RV1 to the skull in huNOG and huNOG-EXL mice. **A,** Bioluminescent images taken of huNOG skulls, **B,** Quantified bioluminescence of huNOG skull metastasis. **C,** Bioluminescent images taken of huNOG-EXL skulls, **D,** Quantified bioluminescence of huNOG-EXL skull metastasis.

**Figure S3:**
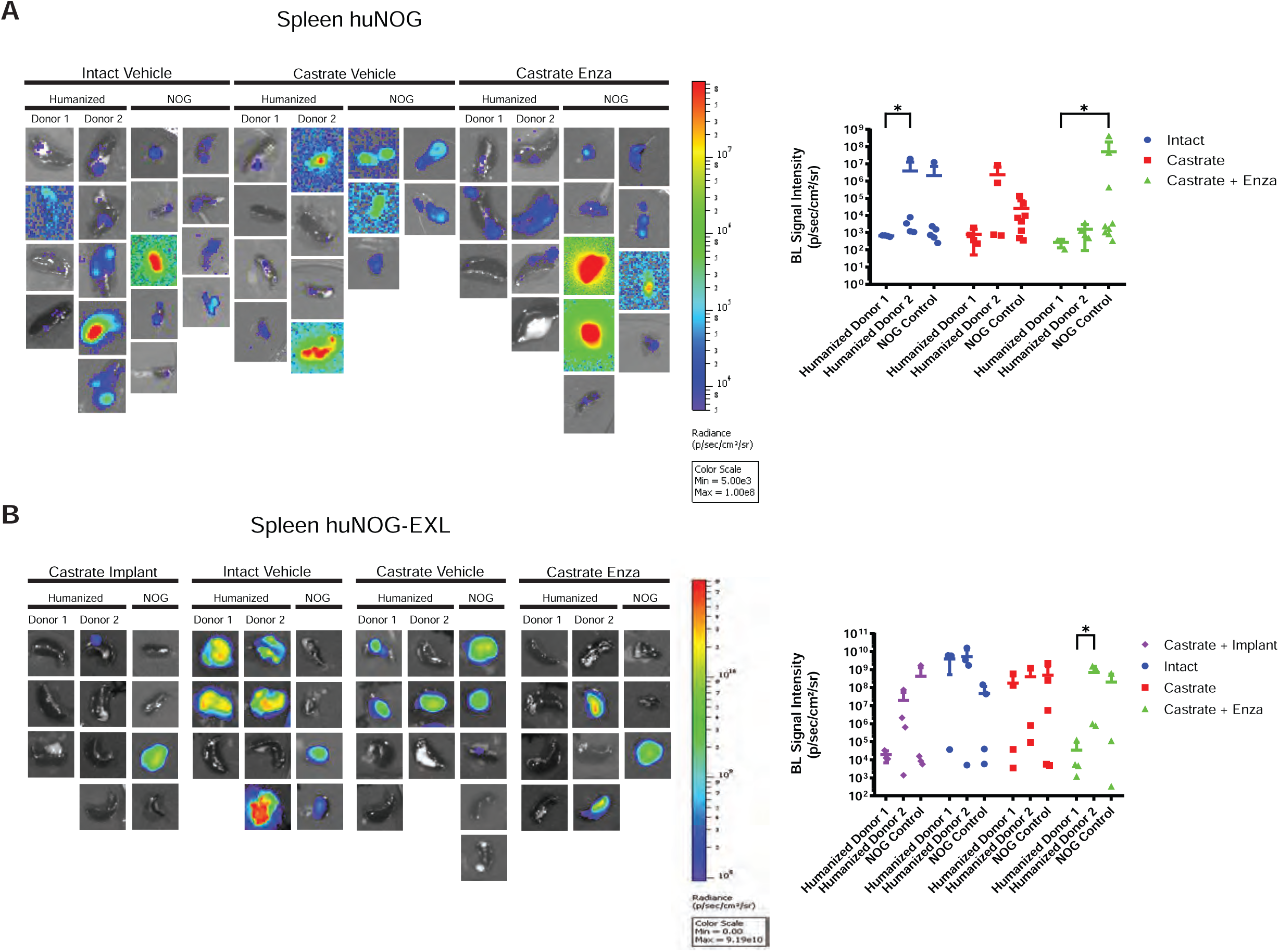
Metastasis by 22RV1 to the spleen in huNOG and huNOG-EXL mice. **A,** Bioluminescent images taken of huNOG spleens, **B,** Quantified bioluminescence of huNOG spleen metastasis. **C,** Bioluminescent images taken of huNOG-EXL spleens, **D,** Quantified bioluminescence of huNOG-EXL spleen metastasis.

**Figure S4:**
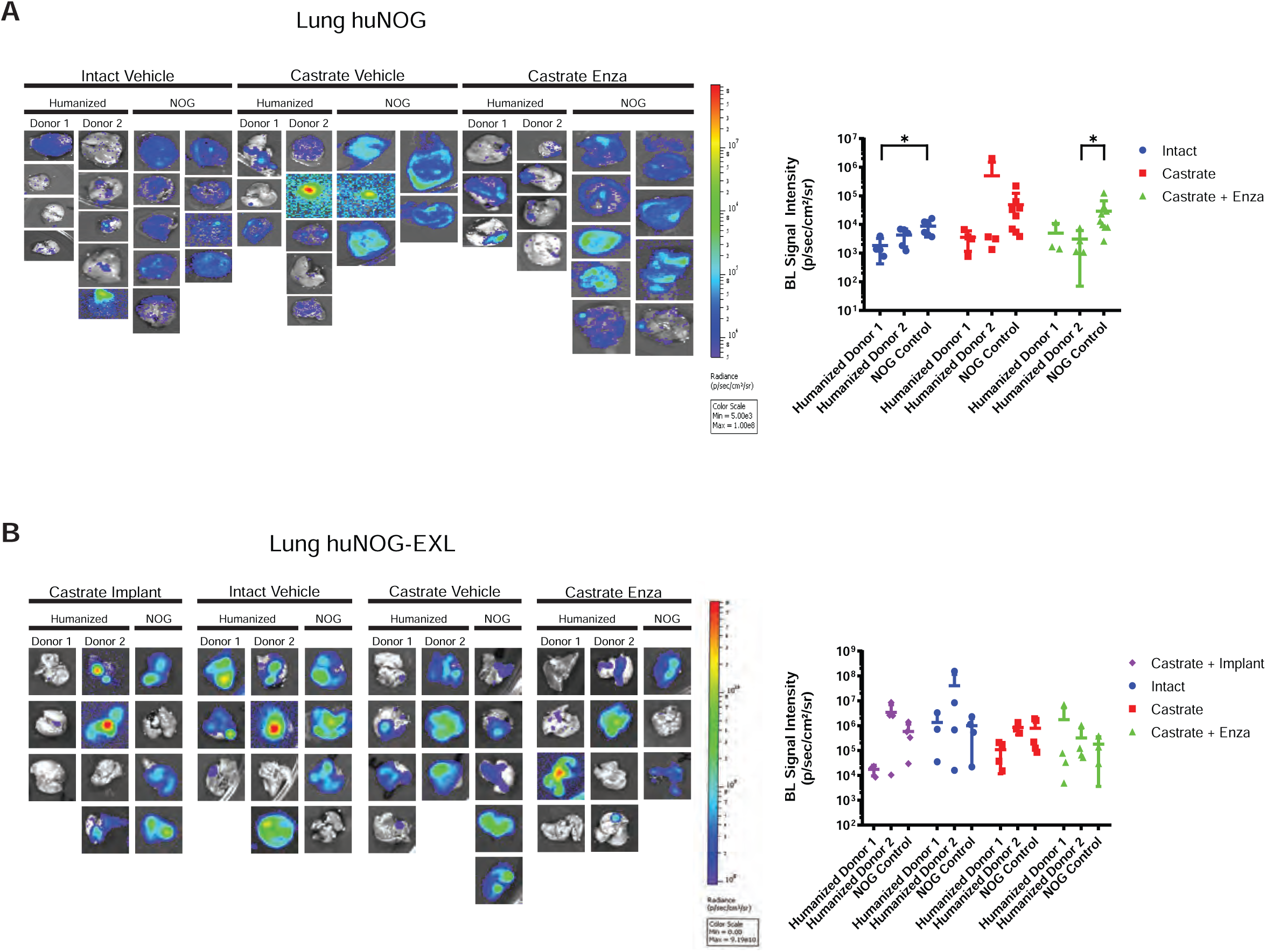
Metastasis by 22RV1 to the lung in huNOG and huNOG-EXL mice. **A,** Bioluminescent images taken of huNOG lungs, **B,** Quantified bioluminescence of huNOG lung metastasis. **C,** Bioluminescent images taken of huNOG-EXL lungs, **D,** Quantified bioluminescence of huNOG-EXL lung metastasis.

**Figure S5:**
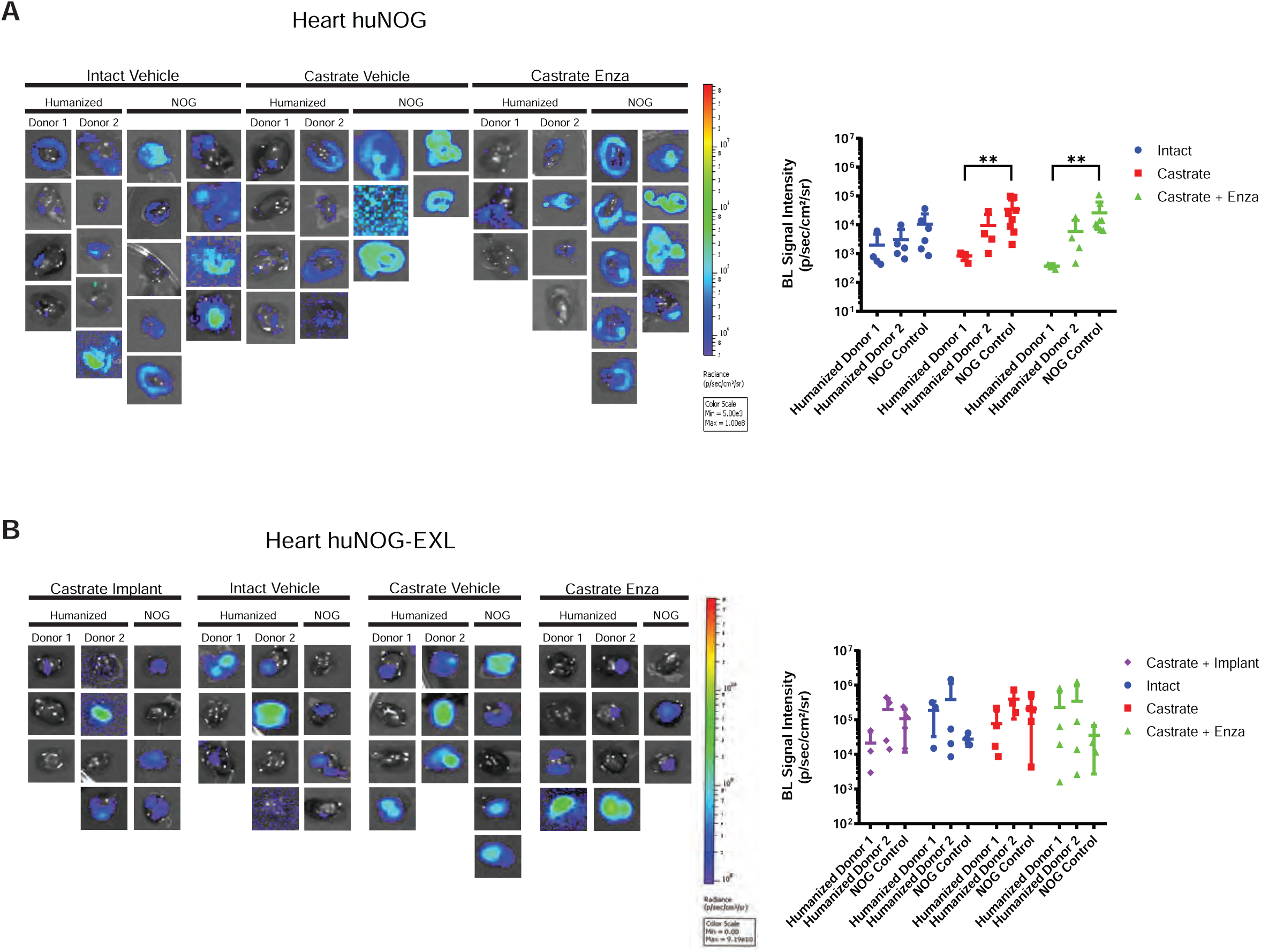
Metastasis by 22RV1 to the heart in huNOG and huNOG-EXL mice. **A,** Bioluminescent images taken of huNOG hearts, **B,** Quantified bioluminescence of huNOG heart metastasis. **C,** Bioluminescent images taken of huNOG-EXL hearts, **D,** Quantified bioluminescence of huNOG-EXL heart metastasis.

**Figure S6:**
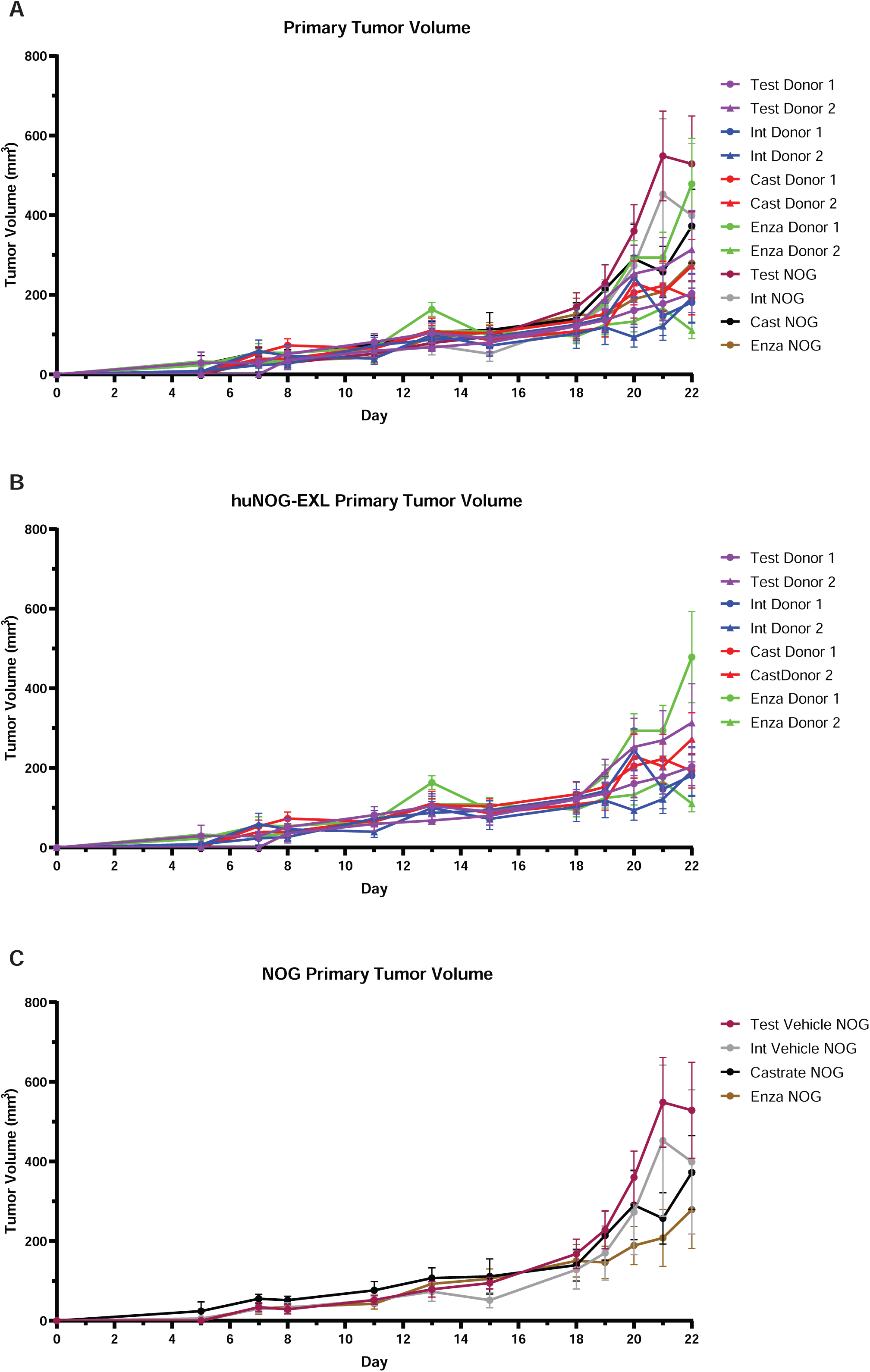
22Rv1 growth in huNOG-EXL and NOG-EXL mice. **A,** Subcutaneous “primary” flank tumor volume growth measured over time in both huNOG-EXL and NOG-EXL mice. **B,** Subcutaneous “primary” flank tumor volume growth measured over time in both huNOG-EXL **C,** Subcutaneous “primary” flank tumor volume growth measured over time in both NOG-EXL.

**Figure S7:**
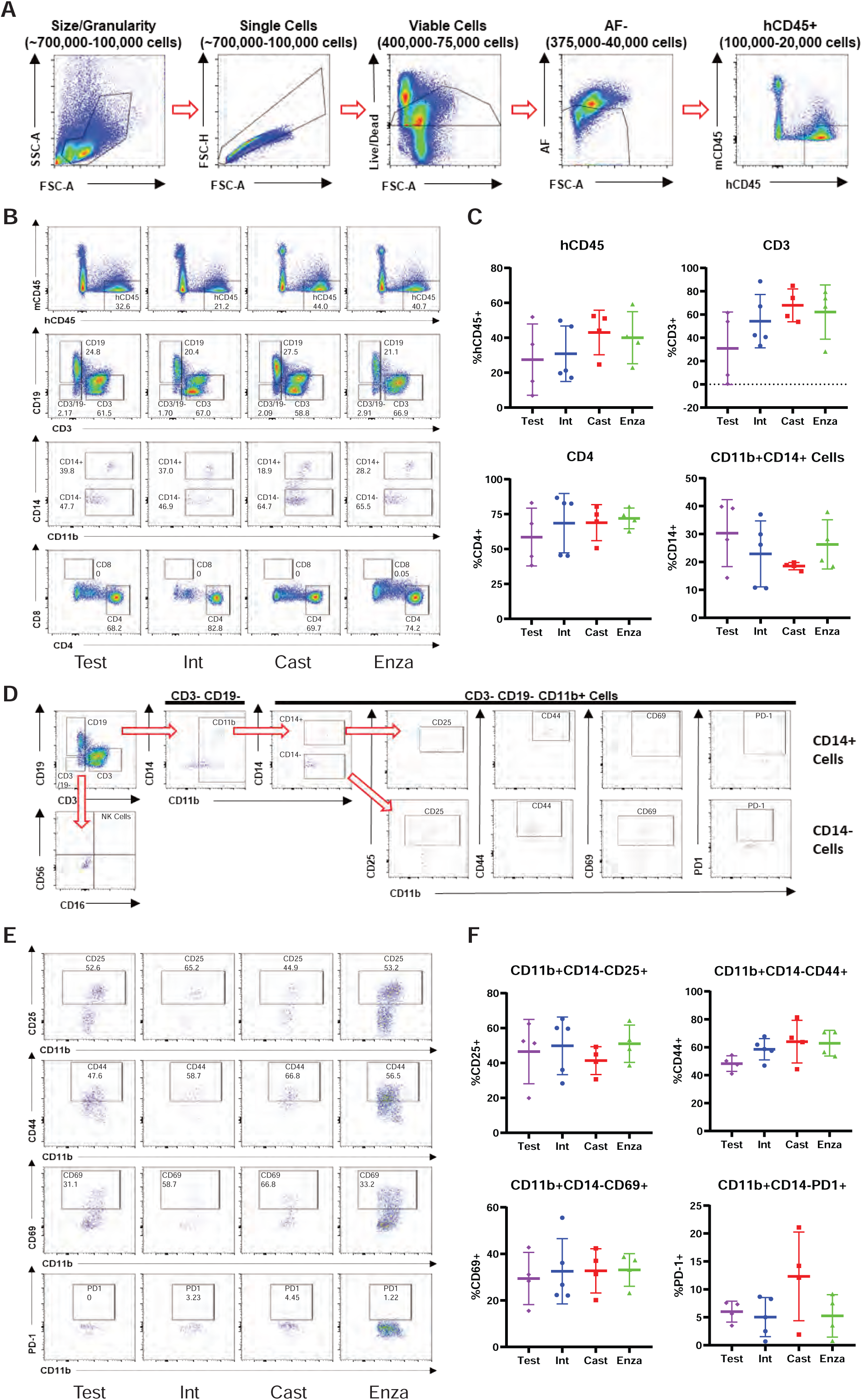
Immune-profile in huNOG-EXL 22Rv1 xenograft spleen. **A,** Gating strategy used to determine presence of human CD45+ cells in the NOG-EXL model spleen. **B,** Representitave data showing the abundance of various immune cell populations; human leukocytes, CD19+ cells, CD3+ cells and double negative cells, MDSC and activated myeloid cells and helper t-cells (CD4+) and cytotoxic t-cells (CD8+) (top to bottom). **C,** Quantitated data comparing human CD45, CD3, CD4, and CD11b populations in spleens isolated from testosterone implanted vehicle, intact vehicle, castrated vehicle, and castrated enzalutamide treated mice. **D,** Gating strategy for determining the activation state of MDSCs (CD3-CD19-CD11b+ CD14-). **E,** Representative data showing the activation state of the MDSC cells harvested from spleens under different treatment categories through the presence of the surface markers: CD25, CD44, CD69 and PD-1. **F,** Quantitated data showing the activation state of the spleen MDSCs throughout the different treatments.

**Figure S8:**
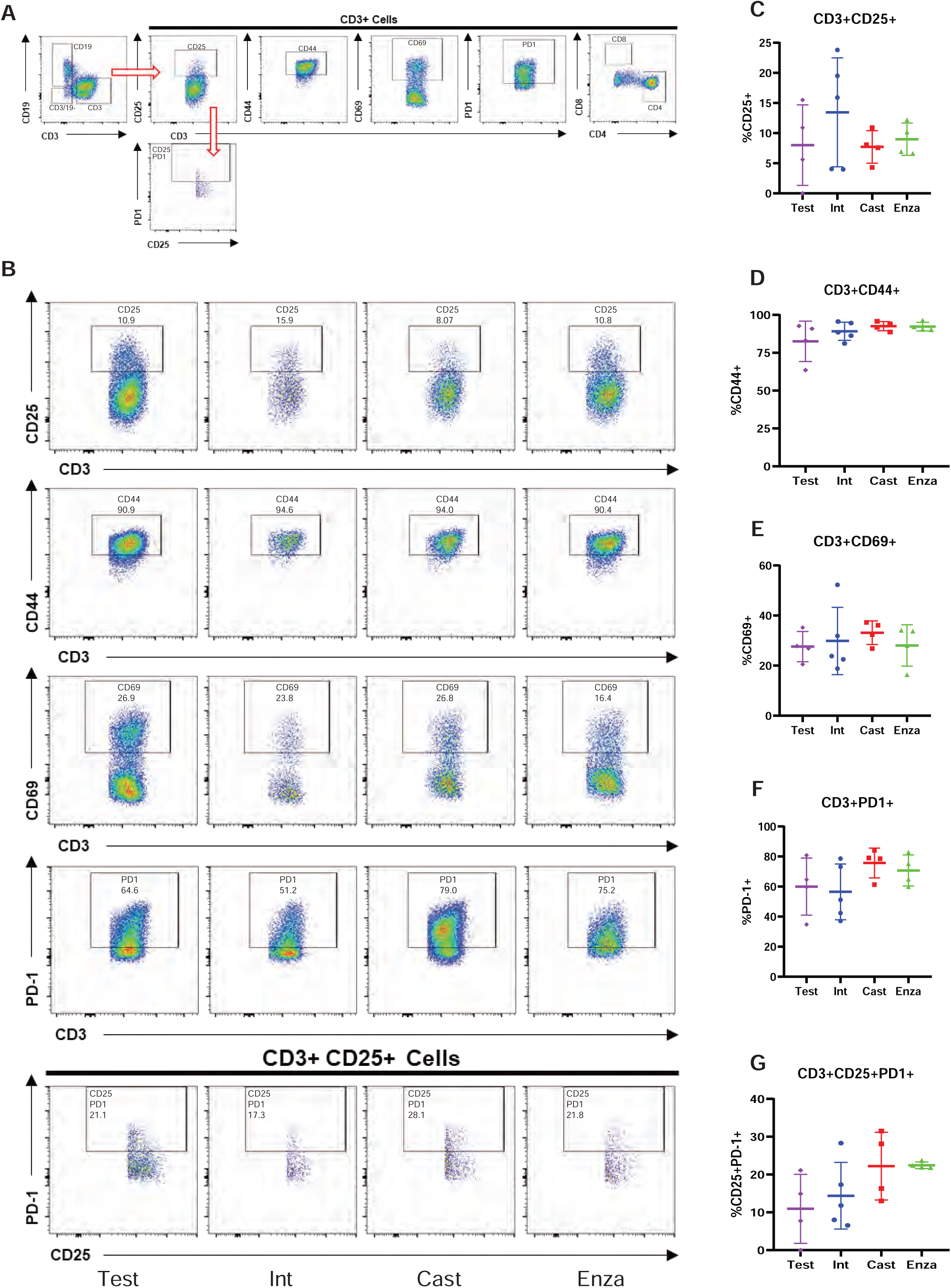
Immune-profile of t-cells in huNOG-EXL spleen. **A,** Gating strategy for determining the activation state of T-cells and regulatory-like T-cells. **B,** Representative data showing the activation state of the CD3+ cells harvested from spleens under different treatment categories through the presence of the surface markers: CD25, CD44, CD69 and PD-1 and regulatory-like cells (CD3+CD25+PD-1+). **C,** Quantitation showing the expression of CD25, **D,** CD44, **E,** CD69, **F,** PD-1, **G,** regulatory-like T-cells (Tregs).

**Figure S9:**
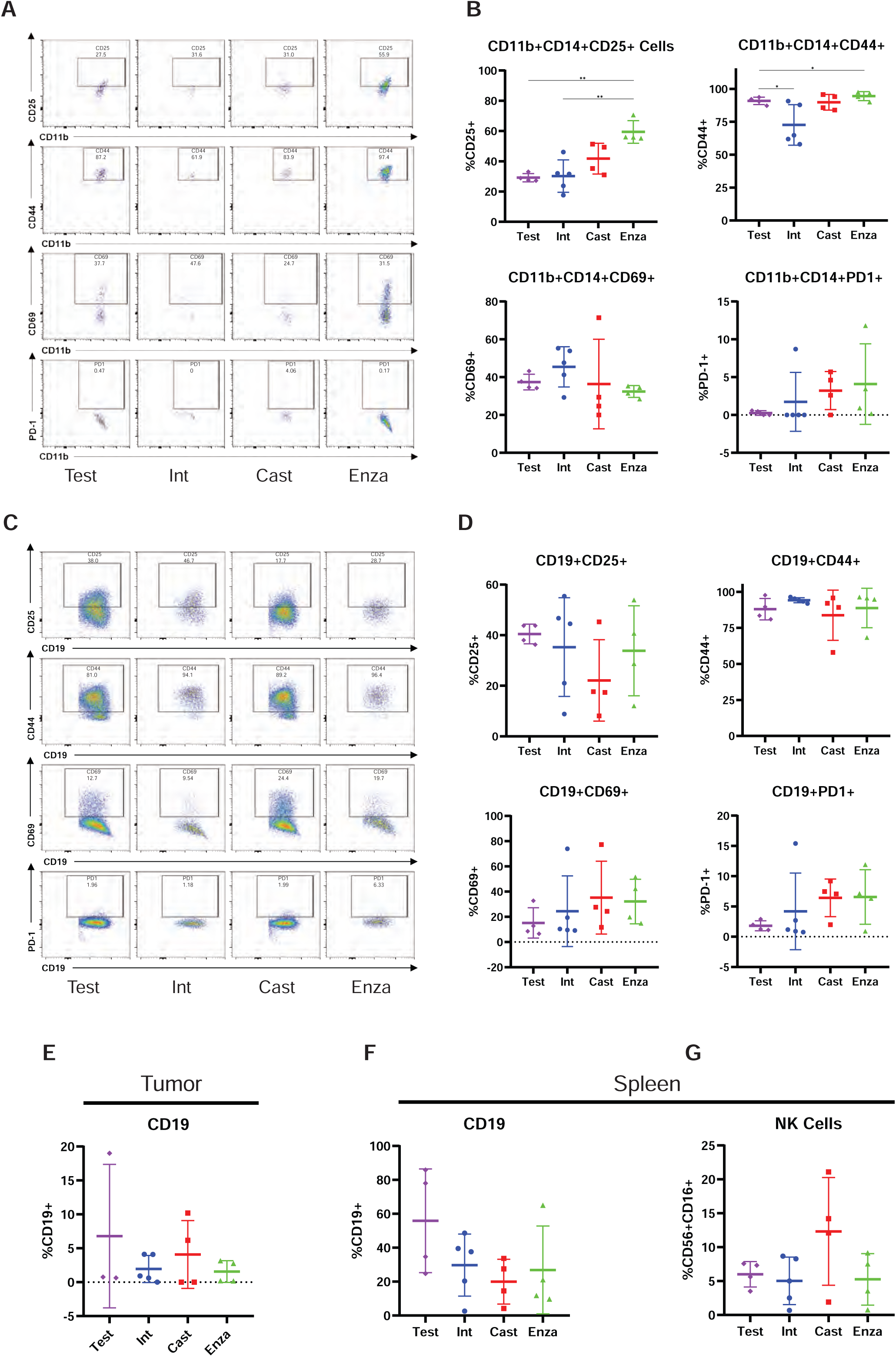
Immune-profile of spleen and tumor B-cell Populations and B-cell activation NK-Cell population in huNOG-EXL spleen. **A,** Representative data showing the activation state of the CD11b+CD14+ cells harvested from spleens under different treatment categories through the presence of the surface markers: CD25, CD44, CD69 and PD-1. **B,** Quantitation showing the expression of CD25, CD44, CD69, PD-1. **C,** Representative data showing the activation state of the CD19+ cells harvested from spleens under different treatment categories through the presence of the surface markers: CD25, CD44, CD69 and PD-1. **D,** Quantitation showing the expression of CD25, CD44, CD69, PD-1. **E,** Percent total CD19+ cells of the total CD45+ population of TILs. **F,** Percent total CD19+ cells of the total CD45+ population of splenocytes. **G,** Percent total NK cells from the spleen (CD3-CD19-CD16+CD56+).

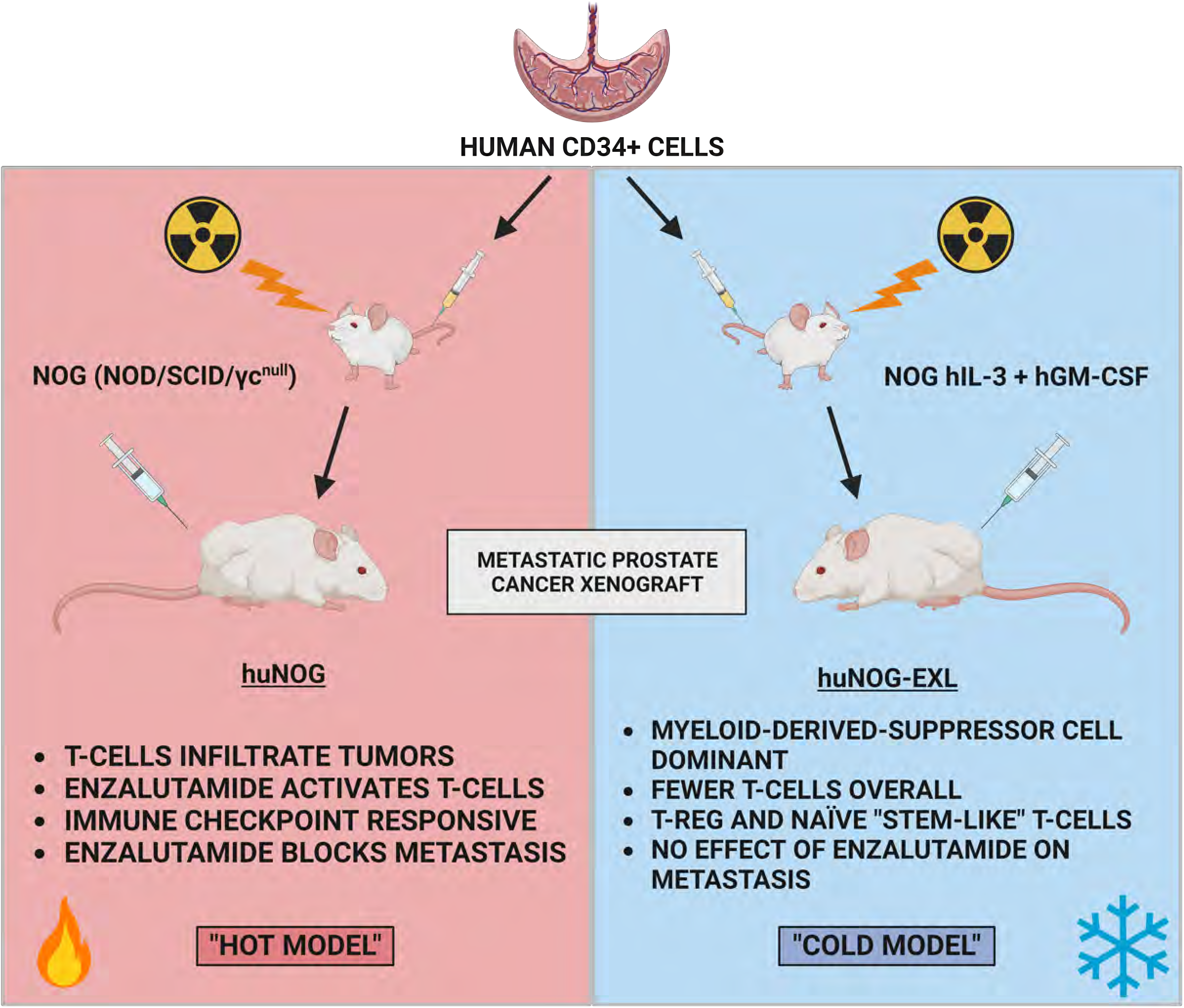

## References

1. He MX, Cuoco MS, Crowdis J, Bosma-Moody A, Zhang Z, Bi K, et al. Transcriptional mediators of treatment resistance in lethal prostate cancer. Nat Med 2021;27:426–33

2. Kregel S, Chen JL, Tom W, Krishnan V, Kach J, Brechka H, et al. Acquired resistance to the second-generation androgen receptor antagonist enzalutamide in castration-resistant prostate cancer. Oncotarget 2016;7:26259–74

3. Ittmann M. Anatomy and Histology of the Human and Murine Prostate. Cold Spring Harb Perspect Med 2018;8

4. Berquin IM, Min Y, Wu R, Wu H, Chen YQ. Expression signature of the mouse prostate. J Biol Chem 2005;280:36442–51

5. Sanmamed MF, Chester C, Melero I, Kohrt H. Defining the optimal murine models to investigate immune checkpoint blockers and their combination with other immunotherapies. Ann Oncol 2016;27:1190–8

6. Ittmann M, Huang J, Radaelli E, Martin P, Signoretti S, Sullivan R, et al. Animal models of human prostate cancer: the consensus report of the New York meeting of the Mouse Models of Human Cancers Consortium Prostate Pathology Committee. Cancer Res 2013;73:2718–36

7. Grabowska MM, DeGraff DJ, Yu X, Jin RJ, Chen Z, Borowsky AD, et al. Mouse models of prostate cancer: picking the best model for the question. Cancer Metastasis Rev 2014;33:377–97

8. Pearson T, Greiner DL, Shultz LD. Creation of “humanized” mice to study human immunity. Curr Protoc Immunol 2008;Chapter 15:15 21 1–15 21

9. Wang Z, Sun K, Xiao Y, Feng B, Mikule K, Ma X, et al. Niraparib activates interferon signaling and potentiates anti-PD-1 antibody efficacy in tumor models. Sci Rep 2019;9:1853

10. Steinkamp MP, O’Mahony OA, Brogley M, Rehman H, Lapensee EW, Dhanasekaran S, et al. Treatment-dependent androgen receptor mutations in prostate cancer exploit multiple mechanisms to evade therapy. Cancer Res 2009;69:4434–42

11. Litvinov IV, Vander Griend DJ, Xu Y, Antony L, Dalrymple SL, Isaacs JT. Low-calcium serum-free defined medium selects for growth of normal prostatic epithelial stem cells. Cancer Res 2006;66:8598–607

12. Kregel S, Kiriluk KJ, Rosen AM, Cai Y, Reyes EE, Otto KB, et al. Sox2 is an androgen receptor-repressed gene that promotes castration-resistant prostate cancer. PLoS One 2013;8:e53701

13. Bluemn EG, Coleman IM, Lucas JM, Coleman RT, Hernandez-Lopez S, Tharakan R, et al. Androgen Receptor Pathway-Independent Prostate Cancer Is Sustained through FGF Signaling. Cancer Cell 2017;32:474–89 e6

14. Kregel S, Bagamasbad P, He S, LaPensee E, Raji Y, Brogley M, et al. Differential modulation of the androgen receptor for prostate cancer therapy depends on the DNA response element. Nucleic Acids Res 2020;48:4741–55

15. Yu J, Green MD, Li S, Sun Y, Journey SN, Choi JE, et al. Liver metastasis restrains immunotherapy efficacy via macrophage-mediated T cell elimination. Nat Med 2021;27:152–64

16. Zhao E, Wang L, Dai J, Kryczek I, Wei S, Vatan L, et al. Regulatory T cells in the bone marrow microenvironment in patients with prostate cancer. Oncoimmunology 2012;1:152–61

17. Ito M, Hiramatsu H, Kobayashi K, Suzue K, Kawahata M, Hioki K, et al. NOD/SCID/gamma(c)(null) mouse: an excellent recipient mouse model for engraftment of human cells. Blood 2002;100:3175–82

18. Li Y, Hwang TH, Oseth LA, Hauge A, Vessella RL, Schmechel SC, et al. AR intragenic deletions linked to androgen receptor splice variant expression and activity in models of prostate cancer progression. Oncogene 2012;31:4759–67

19. Kregel S, Wang C, Han X, Xiao L, Fernandez-Salas E, Bawa P, et al. Androgen receptor degraders overcome common resistance mechanisms developed during prostate cancer treatment. Neoplasia 2020;22:111–9

20. Hussain M, Saad F, Sternberg CN. Enzalutamide in Castration-Resistant Prostate Cancer. N Engl J Med 2018;379:1381

21. Asangani IA, Dommeti VL, Wang X, Malik R, Cieslik M, Yang R, et al. Therapeutic targeting of BET bromodomain proteins in castration-resistant prostate cancer. Nature 2014;510:278–82

22. Hussain M, Fizazi K, Saad F, Rathenborg P, Shore N, Ferreira U, et al. Enzalutamide in Men with Nonmetastatic, Castration-Resistant Prostate Cancer. N Engl J Med 2018;378:2465–74

23. Kazemi MH, Sadri M, Najafi A, Rahimi A, Baghernejadan Z, Khorramdelazad H, et al. Tumor-infiltrating lymphocytes for treatment of solid tumors: It takes two to tango? Front Immunol 2022;13:1018962

24. Brummel K, Eerkens AL, de Bruyn M, Nijman HW. Tumour-infiltrating lymphocytes: from prognosis to treatment selection. Br J Cancer 2023;128:451–8

25. Sieminska I, Baran J. Myeloid-Derived Suppressor Cells as Key Players and Promising Therapy Targets in Prostate Cancer. Front Oncol 2022;12:862416

26. Michiel Sedelaar JP, Dalrymple SS, Isaacs JT. Of mice and men--warning: intact versus castrated adult male mice as xenograft hosts are equivalent to hypogonadal versus abiraterone treated aging human males, respectively. Prostate 2013;73:1316–25

27. Ang JE, Olmos D, de Bono JS. CYP17 blockade by abiraterone: further evidence for frequent continued hormone-dependence in castration-resistant prostate cancer. Br J Cancer 2009;100:671–5

28. Damoiseaux J. The IL-2 - IL-2 receptor pathway in health and disease: The role of the soluble IL-2 receptor. Clin Immunol 2020;218:108515

29. Baaten BJ, Li CR, Bradley LM. Multifaceted regulation of T cells by CD44. Commun Integr Biol 2010;3:508–12

30. Cibrian D, Sanchez-Madrid F. CD69: from activation marker to metabolic gatekeeper. Eur J Immunol 2017;47:946–53

31. Cai J, Wang D, Zhang G, Guo X. The Role Of PD-1/PD-L1 Axis In Treg Development And Function: Implications For Cancer Immunotherapy. Onco Targets Ther 2019;12:8437–45

32. Hirz T, Mei S, Sarkar H, Kfoury Y, Wu S, Verhoeven BM, et al. Dissecting the immune suppressive human prostate tumor microenvironment via integrated single-cell and spatial transcriptomic analyses. Nat Commun 2023;14:663

33. Stultz J, Fong L. How to turn up the heat on the cold immune microenvironment of metastatic prostate cancer. Prostate Cancer Prostatic Dis 2021;24:697–717

34. Sun SH, Benner B, Savardekar H, Lapurga G, Good L, Abood D, et al. Effect of Immune Checkpoint Blockade on Myeloid-Derived Suppressor Cell Populations in Patients With Melanoma. Front Immunol 2021;12:740890

35. Rodems TS, Heninger E, Stahlfeld CN, Gilsdorf CS, Carlson KN, Kircher MR, et al. Reversible epigenetic alterations regulate class I HLA loss in prostate cancer. Commun Biol 2022;5:897

36. Blades RA, Keating PJ, McWilliam LJ, George NJ, Stern PL. Loss of HLA class I expression in prostate cancer: implications for immunotherapy. Urology 1995;46:681–6; discussion 6-7

37. Guan X, Polesso F, Wang C, Sehrawat A, Hawkins RM, Murray SE, et al. Androgen receptor activity in T cells limits checkpoint blockade efficacy. Nature 2022;606:791–6

38. Trigunaite A, Dimo J, Jorgensen TN. Suppressive effects of androgens on the immune system. Cell Immunol 2015;294:87–94

39. Consiglio CR, Udartseva O, Ramsey KD, Bush C, Gollnick SO. Enzalutamide, an Androgen Receptor Antagonist, Enhances Myeloid Cell-Mediated Immune Suppression and Tumor Progression. Cancer Immunol Res 2020;8:1215–27

40. Xu P, Yang JC, Chen B, Nip C, Van Dyke JE, Zhang X, et al. Androgen receptor blockade resistance with enzalutamide in prostate cancer results in immunosuppressive alterations in the tumor immune microenvironment. J Immunother Cancer 2023;11

41. Markowski MC, Shenderov E, Eisenberger MA, Kachhap S, Pardoll DM, Denmeade SR, et al. Extreme responses to immune checkpoint blockade following bipolar androgen therapy and enzalutamide in patients with metastatic castration resistant prostate cancer. Prostate 2020;80:407–11

42. Schweizer MT, Ha G, Gulati R, Brown LC, McKay RR, Dorff T, et al. CDK12-Mutated Prostate Cancer: Clinical Outcomes With Standard Therapies and Immune Checkpoint Blockade. JCO Precis Oncol 2020;4:382–92

